# Post-fire vegetation succession in the Siberian subarctic tundra over 45 years

**DOI:** 10.1101/756163

**Authors:** Ramona J. Heim, Anna Bucharova, Leya Brodt, Johannes Kamp, Daniel Rieker, Andrey V. Soromotin, Andrey Yurtaev, Norbert Hölzel

**Affiliations:** Institute of Landscape Ecology, University of Münster, Heisenbergstraße 2, 48149 Münster, Germany; Research Institute of Ecology and Natural Resources Management, Tyumen State University, 6 Volodarskogo Street, Tyumen, Russia; Department of Conservation Biology, University of Göttingen, Bürgerstr. 50, 37073 Göttingen, Germany; Institute of Ecology, Diversity and Evolution, Goethe University Frankfurt/Main, 60438 Frankfurt am Main, Germany; Institute of Environmental and Agricultural Biology (X-BIO), Tyumen State University, 6 Volodarskogo Street, Tyumen, Russia

**Keywords:** active layer, *Betula nana*, permafrost, plant traits, Russia, soil temperature

## Abstract

Wildfires are relatively rare in subarctic tundra ecosystems, but they can strongly change ecosystem properties. Short-term fire effects on subarctic tundra vegetation are well documented, but long-term vegetation recovery has been studied less. The frequency of tundra fires will increase with climate warming. Understanding the long-term effects of fire is necessary to predict future ecosystem changes.

We used a space-for-time approach to assess vegetation recovery after fire over more than four decades. We studied soil and vegetation patterns on three large fire scars (>44, 28 and 12 years old) in dry, lichen-dominated forest tundra in Western Siberia. On 60 plots, we determined soil temperature and permafrost thaw depth, sampled vegetation and measured plant functional traits. We assessed trends in NDVI to support the field-based results on vegetation recovery.

Soil temperature, permafrost thaw depth and total vegetation cover had recovered to pre-fire levels after >44 years, as well as total vegetation cover. In contrast, after >44 years, functional groups had not recovered to the pre-fire state. Burnt areas had lower lichen and higher bryophyte and shrub cover. The dominating shrub species, *Betula nana*, exhibited a higher vitality (higher specific leaf area and plant height) on burnt compared with control plots, suggesting a fire legacy effect in shrub growth. Our results confirm patterns of shrub encroachment after fire that were detected before in other parts of the Arctic and Subarctic. In the so far poorly studied Western Siberian forest tundra we demonstrate for the first time, long-term fire-legacies on the functional composition of relatively dry shrub- and lichen-dominated vegetation.

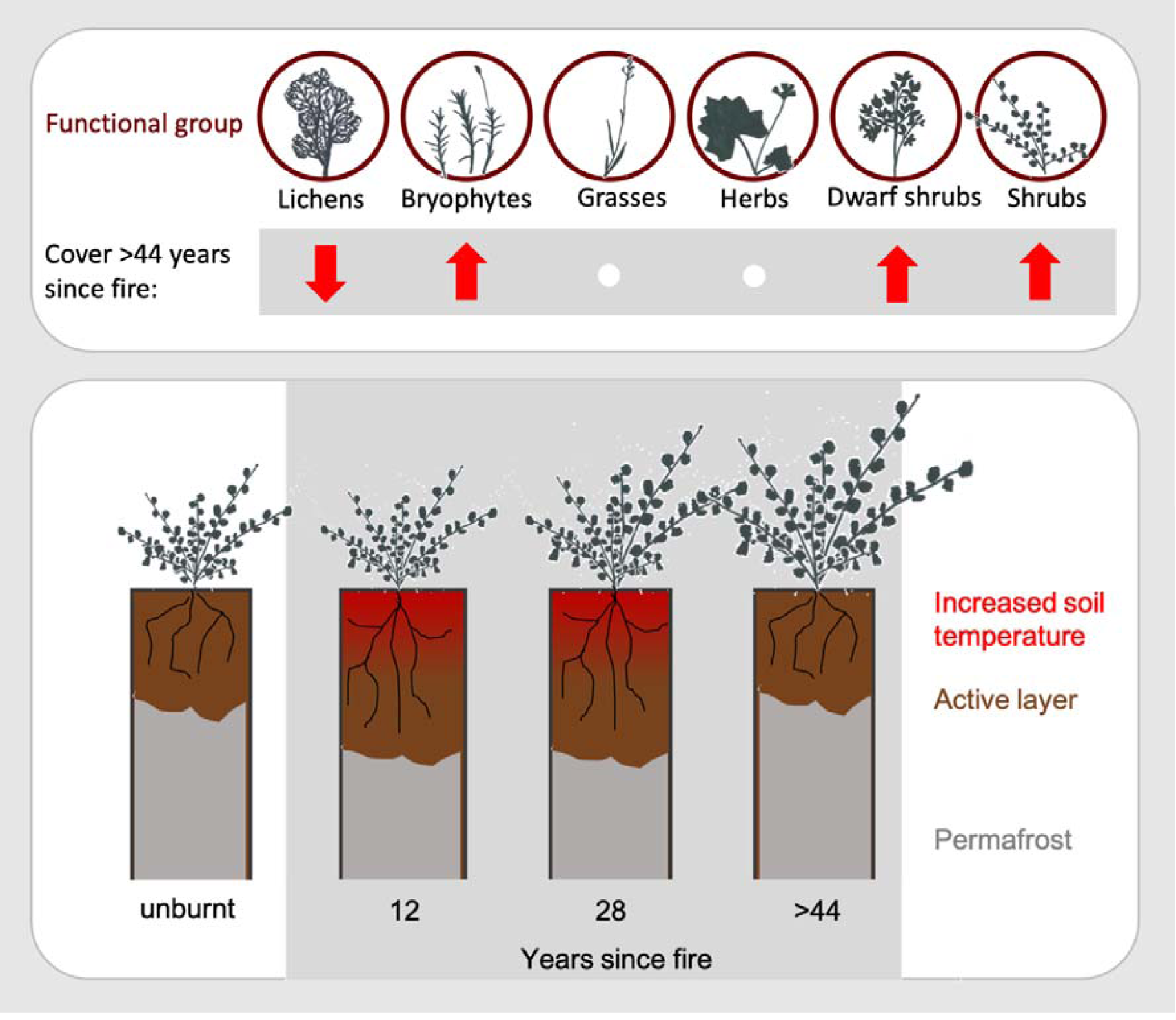

## 1. Introduction

Fire drives ecosystem patterns and processes across biomes and climatic zones (Bowman et al. 2009). In arctic and subarctic tundra, fires are naturally rare, cool and small compared to other biomes (Viereck and Schandelmeier 1980, Archibald et al. 2013), but their frequency and extent is predicted to increase with climate change (Flannigan et al. 2009). This will affect ecosystem functioning (Hu et al. 2015), and may change the tundra biome from a carbon sink to a carbon source (Abbott et al. 2016).

Large tundra fires occur during extended warm periods without precipitation when vegetation and litter are dry (Hu et al. 2015, Masrur et al. 2018). With climate change, summers are expected to become drier, and the duration of the vegetation period will increase leading to a higher ignition probability and a larger size and intensity of tundra fires (Moritz et al. 2012, Young et al. 2016).

Disturbance by fire changes vegetation composition and structure and influences soil properties of the tundra (Narita et al. 2015, Mekonnen et al. 2019). Vegetation is directly affected by combustion of living and dead biomass (litter). After severe fires, vegetation cover is reduced, surface albedo decreases, more solar energy is transferred to the ground and soil temperatures increase (Rocha and Shaver 2011, French et al. 2016). Aboveground biomass insulates soil against extreme temperatures and thereby buffers temperature fluctuations. Consequently, the loss of vegetation can result in higher summer soil temperatures, causing permafrost thaw and a subsequent increase in permafrost thaw depth (Chambers et al. 2005, Jiang et al. 2015a, Zhang et al. 2015). Increased soil temperature and subsequent permafrost thaw depth lead to a higher nitrogen availability to plants through thawing and enhanced microbial activity (Keuper et al. 2012). Further nutrients, mainly inorganic nitrogen, are provided by remaining ash after fire (Jiang et al. 2015b).

Regeneration patterns and processes after fire differs between vascular plants and cryptogams (Bret-Harte et al. 2013). Lichen and moss recovery after fire is very slow, probably because cryptogam biomass is completely destroyed during fire and burnt areas need to be recolonized, while shrubs and graminoids, which often benefit from fire, can regenerate from persistent bud and seed banks (Racine et al. 2004, Jandt et al. 2008, Turetsky et al. 2012, Narita et al. 2015). A depleted lichen cover after fire is often regarded as a problem for reindeer grazing, as animals do not find enough food during winter if extended areas of their forage range burnt (Viereck and Schandelmeier 1980). A general decline in lichens represents a threat for indigenous people who depend on reindeer herds (Sandström et al. 2016). Fire amplifies the shrub expansion that has been observed in many regions across the tundra, and has been attributed to global warming (e.g. Arefiev 2016, McLaren et al. 2017). Shrub encroachment leads to increased shading of the soil surface in summer and protects permafrost from thawing (Myers-Smith et al. 2011), but can also reduce permafrost depth, because shrubs retain more snow during winter, which insulates the soil against frost (Myers-Smith and Hik 2013). Thicker snow layers result in a higher amount of meltwater in summer that increases thermal conductivity, leads to soil warming in deeper layers and ultimately thawing permafrost (Johansson et al. 2013). Moreover, shrubs have a low albedo, warm up in the sun above the snow layer and thus accelerate snow melt in spring (Loranty et al. 2018).

While it is known that fire promotes the growth of shrubs and graminoids, we know less about the mechanisms behind this pattern of enhanced growth and how these change over time. Plant functional traits are established ecosystem properties that link plant physiology and environmental processes and therefore provide valuable information about post-fire succession (Keeley et al. 2011, Cornelissen and Makoto 2014). Tundra fires alter surface energy dynamics and nutrient availability through changes in vegetation and soil properties (Keuper et al. 2012, Loranty et al. 2018), and some functional traits capture these chances, especially plant height, specific leaf area and decreases leaf dry matter content (Hobbie et al. 2002, Bjorkman et al. 2018). Shifts in plant functional traits along post-fire successional gradients will provide insight how the post-fire environment changes affect plants.

Most of our knowledge on the effect of fire on tundra ecosystems comes from Alaska (Racine et al. 1987, e.g. Barrett et al. 2012, Bret-Harte et al. 2013). However, Alaska comprises less than 10% of global Arctic vegetation (Walker et al. 2005). Considerably less is known about other parts of the world, particularly about the vast areas of the Siberian tundra (Frost and Epstein 2014, Loranty et al. 2014, e.g. Abdulmanova and Ektova 2015). Although the tundra ecosystems in Alaska and Russia are floristically similar (Walker et al. 2005), their ecological history significantly differs. In contrast to Alaska, where herds of wild caribous can be found, the north of Western Siberia has been influenced by reindeer pastoralism for centuries (Forbes and Kumpula 2009), which has a strong impact (e.g. defoliation, nutrient availability) on tundra vegetation (Sundqvist et al. 2019). Moreover, most fire studies in Alaska were conducted in relatively humid climatic and edaphic conditions with predominantly moist tussock graminoid-dominated tundra (Racine et al. 1987, Mack et al. 2011, Narita et al. 2015). Considerably less is known about post-fire recovery over time of dry, lichen-dominated tundra vegetation, common in Siberia, and the effects of those shifts on ecosystem properties. As lichens and tussock graminoids respond in different ways to fire, it is likely that the two vegetation types will differ in post-fire succession (Jandt et al. 2008). Understanding the post-fire recovery of Siberian dry forest tundra is essential for a comprehensive assessment of fire effect and post-fire regeneration of global arctic ecosystems.

Here we present the first comprehensive study on long-term effects of fires on dry forest tundra vegetation at the northern edge of forest tundra ecozone of Western Siberia. Using a space-for-time approach, to extrapolate the temporal trend of recovery after fire from fire scars of different age while assuming spatial and temporal variation to be similar, we studied post-fire vegetation succession and soil characteristics on three large fire scars (>44, 28 and 12 years old). The goal of our study was to assess long-term effects of tundra fire on vegetation and in detail we anticipated that:

1. On burned plots, the cover and height of vascular plants was higher compared to unburned plots while the cover and height of lichens and bryophytes was lower.
2. Fire-related changes in vegetation structure affected soil temperature and the depth of the permafrost thaw.
3. Fire-related changes in soil temperature led to higher values in growth-related plant functional traits in vascular plants.

## 2. Material and Methods

### 2.1. Study area

Our study area is situated on the northern border of the forest tundra ecozone in Western Siberia within the Yamalo-Nenets Autonomous Okrug between the rivers Pur and Taz, north of the Arctic Circle (centre of the study area at 67° 1’19.59”N, 79° 1’53.53”E, total study area size ca. 70 km^2^, see Fig. 1).

**Fig. 1:**
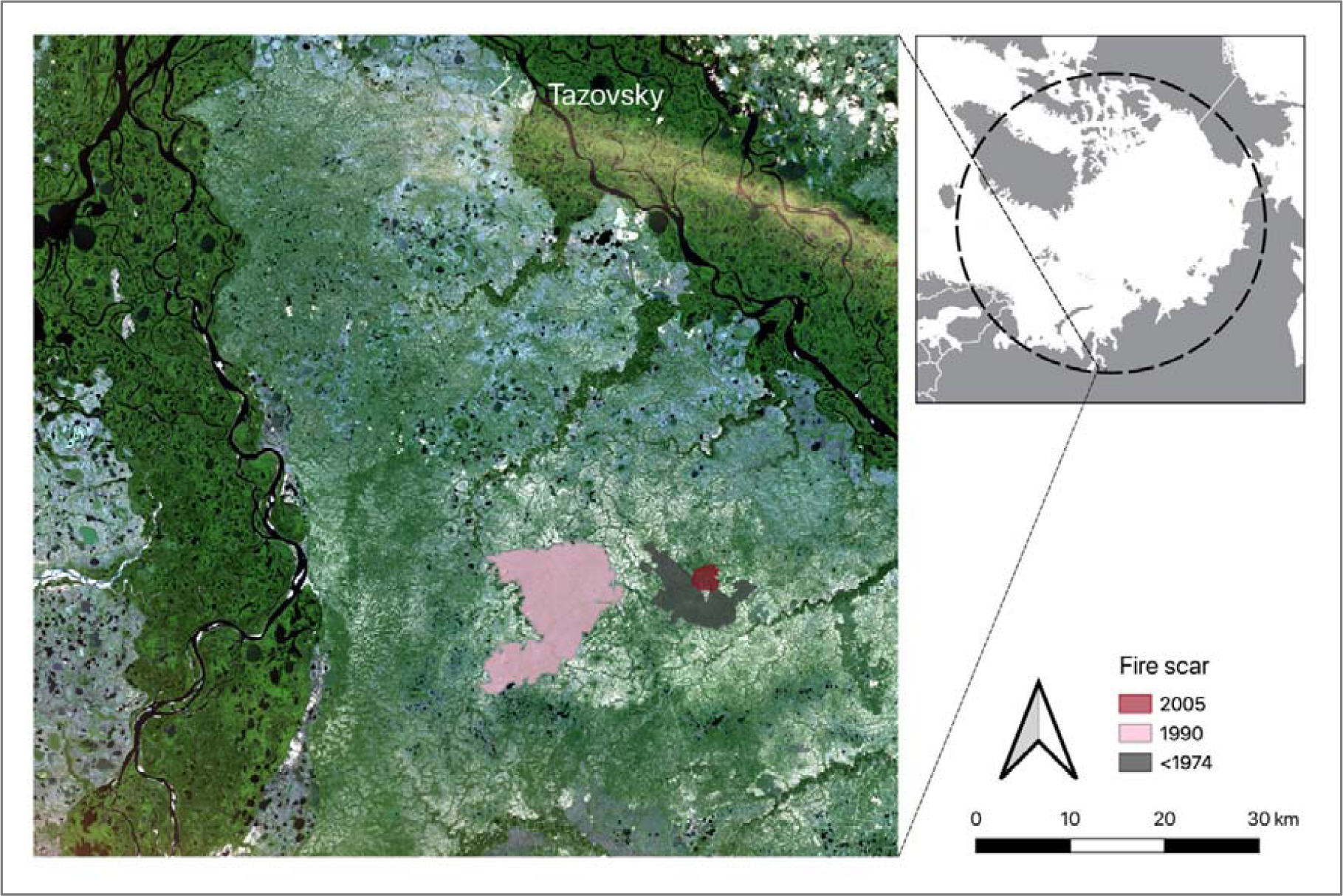
Study area with three fire scars in the north of Western Siberia. Satellite image source: Copernicus Sentinel-2 data (2019).

The region has a subarctic climate with a mean annual air temperature of −8.1 °C, a mean January air temperature of −26.2 °C and a mean July air temperature of 14.4 °C (Kazakov 2019). Annual precipitation amounts to 482 mm with a maximum in August. The growing season lasts from mid-June to early September (Kazakov 2019). The study area is located in the ecotonal transition between the forest tundra to the south and the open treeless shrub tundra to the north (Yurkovskaya 2011). Vegetation is dominated by reindeer lichens (mostly *Cladonia stellaris* and *Cladonia rangiferina*) that cover in thick layers around ca. 70% of the soil surface, shrubs such as *Betula nana* (ca. 25% cover), and dwarf shrubs such as *Vaccinium uliginosum* (ca. 10 % cover) (Suppl. A1). Herbs and graminoids as well as bryophytes are far less abundant. The largely open landscape is sparsely dotted with larch trees (*Larix sibirica*), the only tree species occurring in our study area.

With the exception of depressions and small streams, the study region is well-drained in comparison with other tundra areas e.g. relatively moist tundra types in Alaska. Soils are Cryosols (IUSS Working Group WRB 2015) that developed in silty, loess-like parent material. Soil organic layer thickness was generally low (mean of 4 cm), on all burnt and unburnt sites. Mean topsoil pH (0-5 cm depth) had a mean of 4.3 and mean topsoil concentrations of Carbon and Nitrogen were 0.82 % and 0.06 %, respectively. Fires in the region are mainly caused by lightning (Kornienko 2018), but with a recent increase in transport and settlement infrastructure in the area (due to oil and gas exploitation) the number of human-induced fires has increased (Vilchek and Bykova 1992, Mollicone et al. 2006, Yu et al. 2015).

From 1985-2017, 10.5% of the forest tundra of Western Siberia was affected by fires, most of them occurring in the vegetation type of dwarf shrub-, lichen- and moss-dominated forest tundra with larch trees (Moskovchenko et al. 2020). Frequency and intensity of fires, occurring in this region are varying strongly as well as the area burnt, dependent on weather conditions (Moskovchenko et al. 2020). Summer temperatures and precipitation are linked to the burnt area and thus most fires occur, when flammable material is dry (Moskovchenko et al. 2020).

### 2.2. Field sampling

We compared vegetation and soil parameters at three large, non-overlapping burnt areas (fire scars) and adjacent unburnt control sites. All three sites were located close to each other (< 10 km distance between scars) in an area with homogenous environmental conditions in terms of parent rock and topography.

Fire scars were detected by comparing annual Landsat images in Google Earth Timelapse visually (Gorelick et al. 2017) back to the year 1985, and by visually inspecting older Landsat images back to 1973 that were downloaded via USGS earth explorer (U.S. Geological Survey 2018). The RGB band combination was set to 7,5,4, to make the spectral appearance comparable to the images displayed in the timelapse tool. Pale areas were considered unburnt as they had a high lichen cover in the year before the fire. Areas that switched from pale to dark were identified as fire scars. They were readily distinguished from other features with similar spectral properties by their irregular shape, and the location of their borders that generally followed meandering brooks. The youngest fire scar (542 ha) burnt in 2005, a medium-aged scar (ca. 12,500 ha) burnt in 1990, and an old scar (3,500 ha) was already present on the first satellite image we used (Landsat 1) from 1973. In the field, the burn status of selected plots was checked again using signs of burnt trees and charcoal on the ground.

The fire severity at the youngest and the medium-aged fire scar did not differ significantly (see Suppl. A2). A common tool to estimate fire severity is the Normalized Burn Ratio Index (NBR), which is calculated by using near-infrared and shortwave-infrared wavelength (Key and Benson 2006). A large difference between pre- and postfire Normalized Burn Ratio Index (dNBR) indicates high fire severity. The mean dNBR for all burnt plots of the youngest fire scar was moderate to high with dNBR = 0.56 (dNBR_range_ = 0.36-0.68). Similar results were found for the medium-aged fire scar with dNBR = 0.60 (dNBR_range_ = 0.37-0.71) (Key and Benson 2006). We could not evaluate fire severity for the older fire scar because of the lack of pre-fire satellite images.

Fieldwork took place in July 2017 (at the areas burnt before 1973 and in 2005) and July 2018 (at the area burnt in 1990). Time since fire was therefore 12, 28 and at least 44 years for the three scars. As the tundra ecosystem is rather stable with low inter-annual variability in environmental conditions (Dahl 1975), sampling in two subsequent years is unlikely to bias the results, especially as the climatic conditions in the two years were rather similar. Mean temperature in the years 2016, 2017 and 2018 were −4.8, −5.6 °C and −6.7°C, respectively. Warmest month was July in both years with a mean of 18.8 °C in 2018, 15.8° C in 2017 and 14.1 °C in 2018. Coldest month was December 2016 (−29.7 °C), January 2017 (−25.7 °C) and February 2018 (−23.2 °C). Maximum snow height was 81 cm in 2016, 91 cm in 2017 and 107 cm in 2018. In 2016 there was snow on 219 days and in both subsequent years, there was snow on 237 days (Kazakov 2019).

On each of the three fire scars, we selected 10 sampling locations along the fire border, which were located at least 300 m from each other to cover local environmental variation. At each location, we recorded vegetation and environmental parameters at one burnt and one unburnt plot of 10 × 10 m, resulting in a total of 60 plots. The plots of each pair were placed as close to each other as possible, but at least 100 m apart. Minimum plot distance from the fire scar border was always 50 m to avoid edge effects.

On each plot, we visually estimated total vegetation cover, bare soil and the cover and height of different functional plant groups (lichens, bryophytes, herbs, graminoids, dwarf shrubs, shrubs) (Suppl. A3). Vegetation cover was visually estimated in % on 1m × 1m plots and extrapolated to the 10m × 10m plots. For the most abundant dwarf shrub and shrub species, *Vaccinium uliginosum* L. and *Betula nana* L., we measured six functional plant traits that are related to plant growth (Hudson et al. 2011, Perez-Harguindeguy et al. 2016, Bjorkman et al. 2018): Mean aboveground cover for plant individuals (calculated as: (*longest diameter + diameter orthogonal to longest diameter) / 2*), plant height, blade length, leaf thickness, specific leaf area (SLA) and leaf dry matter content (LDMC). The traits were measured on one mature, healthy individual per species and plot. Apart from dry weights for SLA and LDMC, all measures were taken in the field. To obtain leaf area for the SLA, we placed at least four leaves per plant on an illuminated plate equipped with calibration scale and took photographs in the field. The leaves were dried, stored and weighted in the laboratory. Leaf thickness was measured using a digital thickness gauge.

On all plots, we measured soil temperature 12 cm below the soil surface and permafrost thaw depth using a thermometer and a metal stick. Both measurements were repeated five times per plot (in each of the four corners and the centre) and the mean was calculated.

### 2.3. NDVI change analysis

To complement the field analysis, an NDVI value characterizing each sample plot was derived from Landsat images for all burnt and unburnt plots of the 1990 fire scar (as this was the oldest scar for which satellite images before the fire event were available). NDVI was calculated for each vegetation sampling plot inside and outside the burnt area of 1990 (choosing the pixel, which included the plot coordinates). We derived NDVI data from 16 different scenes of Landsat 5, 7 and 8 (spatial resolution of 30 m per pixel) from mid-July 1990 (before the fire event) to 2017. The Landsat scenes were downloaded via the USGS earth explorer (U.S. Geological Survey 2018). Due to the short growing season in the study region, we included only images of July and August in our analysis. We processed satellite images performing atmospheric correction and reduced the differences between satellite systems using the FLAASH module in ENVI 4.8 (ITTVIS) and combined channels (3, 2 and 1 for Landsat 5 and 7 and 4, 3 and 2 for Landsat 8). We calculated the vegetation index in ENVI 4.8 (ITTVIS) according to the formula: *NDVI = (NIR – RED) / (NIR + RED)*, where NIR is the reflection in the near infrared region of the spectrum (0.7-1.0 μm), and RED the reflection in the red region of the spectrum (0.6-0.7 microns).

### 2.4. Statistical analysis

To test the anticipation 1 (on burned plots, the cover and height of vascular plants was higher compared to unburned plots while the cover and height of lichens and bryophytes was lower) we subtracted the cover and height values of the control plots from those of the paired plots within the burnt area to obtain a difference. We then modelled the difference in vegetation cover and height between fire and control plots as a function of the vegetation recovery time (years since fire event) using linear mixed-effects models in a Bayesian framework in R, Version 3.5.3 (R Core Team 2019). Difference in cover and height were the response variables. Functional vegetation group and years since fire, and their interactions were fitted as fixed effects. Plot ID was included as a random effect to account for the repeated sampling of vegetation groups on each plot. We used the R-package lme4 (Bates et al. 2015). In the model with cover as dependent variable, the variance of the random factor was 0. Therefore, we excluded the random factor “plot ID”.

In order to obtain the posterior distribution, we used improper priors and simulated 2000 values from the joint posterior distributions of the model parameters using the sim function from the R-packages arm (Gelman and Su 2007) and blmeco (Korner-Nievergelt et al. 2015). The posterior distribution is a probability distribution that summarises updated beliefs about the parameter after observing the data and is thus a combination of the prior distribution and the likelihood function. We present the mean values and the 95% credible interval (CrI) of the simulated posterior distribution. The 95% CrI represents the range within the true value is expected with a probability of 0.95 and is limited by the 2.5% and the 97.5% quantile of the posterior distribution. We calculated the posterior probabilities for anticipation 1 by using the proportion of simulated values of the posterior distribution being > 0. A posterior probability of 1 means that all fitted values are larger than 0. Regarding our anticipation this would mean that the burnt plots have a significantly higher cover than unburnt plots. A posterior probability of 0 means the opposite: all fitted values are smaller than 0. A posterior probability of 0.5 indicates that means of the distributions of fitted values are identical to 0 and burnt and unburnt plots do not differ.

For testing anticipation 2 (fire-related changes in vegetation structure affected soil temperature and the depth of the permafrost thaw) we subtracted the soil temperature and permafrost thaw depth values of the control plots from those of the paired plots within the burnt area. We then fitted linear mixed-effects models with difference in soil temperature and difference in permafrost thaw depth as response variables and years since fire event as an independent variable. Plot was fitted as random effect. We calculated the posterior probabilities of anticipation 2 as described above in the analysis for anticipation 1.

We furthermore subtracted total cover and bare soil values of the control plots from those of the paired plots within the burnt area and fitted again linear mixed-effects models with difference in soil temperature and difference in permafrost thaw depth as response variables and years since fire event as an independent variable. Plot was fitted as random effect, but excluded from the final model, as it explained no variance. We calculated the posterior probabilities as described above.

Anticipation 3 (fire-related changes in soil temperature led to higher values in growth-related plant functional traits in vascular plants) was tested by modelling the differences in six different plant traits between the burnt and control plot as a function of time since fire, for *V. uliginosum* and *B. nana*. We used linear mixed-effects models with difference in trait value (traits: mean aboveground cover, plant height, blade length, leaf thickness, SLA, LDMC) as dependent variable, species, years since fire event and their interaction as fixed effects, and plot as random effect. The posterior probabilities were calculated as described before. For testing if the found differences between burnt and unburnt plots decreased during recovery, we repeated the analysis that was used to test anticipation 1 and 3 but with the years since fire as a continuous covariate. We calculated the posterior probabilities of the model used to test anticipation 4 by using the proportion of simulated values of the posterior distribution for the slopes being > 0. We could not date the oldest fire event exactly as no satellite images from the years prior to 1973 were available. We therefore performed a sensitivity analysis. We re-run the model to test anticipation 4 (Scenario1) assuming that the oldest fire scar burnt 100 years ago (instead of 44 years as in the models above) (Scenario2) and compared the results.

As complementation to the field data, we modelled the NDVI difference between burnt and reference plot as a function of time since fire using a linear model (LM) with a quadratic term. The posterior probabilities were calculated as described before.

Model assumptions for all described analyses were graphically assessed with Tukey-Anscombe- and QQ-plots. Figures were created using packages ggplot2 (Wickham 2009) and cowplot (Wilke 2019). Goodness of fit was determined with R^2^ using the MuMIn-package (Barton 2019).

## 3. Results

### 3.1. Influence of fire on soil temperature and permafrost thaw depth

Fire strongly influenced soil temperature and permafrost thaw depth. Twelve and 28 years after the event, burnt plots showed a higher soil temperature and a deeper permafrost thaw than control plots (Fig. 2, Table 1, Suppl. A4). Four decades after fire, the differences were no longer apparent, indicating that soil temperatures and permafrost thaw depth returned to levels of unburnt plots.

**Table 1:** Mean values of change in soil temperature (12 cm depth) and permafrost thaw depth, estimated by linear mixed-effects models including years since fire and mean values of change in total cover and bare soil, estimated by a linear mixed-effects model, including years since fire and the different groups. Means, 2.5%, 97.5% quantiles of the posterior distributions are given. Effects of change are shown in bold, if there is a high probability of the estimate to be different from 0 (P(*estimate*>0)>0.95, P(*estimate*>0)<0.05).

| Variable | Years since fire | Mean | 2.50% | 97.50% | $P(\text{estimate} > 0)$ |
| --- | --- | --- | --- | --- | --- |
| *Soil temperature*<br>$R^2 = 0.53$ | **12** | **6.29** | **4.61** | **8.19** | **1.00** |
|  | **28** | **6.29** | **4.48** | **8.02** | **1.00** |
|  | >44 | 0.24 | -1.52 | 1.97 | 0.62 |
| *Active layer depth*<br>$R^2 = 0.55$ | **12** | **-33.02** | **-44.53** | **-20.66** | **0.00** |
|  | **28** | **-32.20** | **-43.61** | **-20.58** | **0.00** |
|  | >44 | -6.62 | -5.18 | 17.74 | 0.88 |
| *Total cover*<br>$R^2 = 0.71$ | **12** | **-27.40** | **-32.41** | **-22.48** | **0.00** |
|  | **28** | **-14.65** | **-19.58** | **-9.87** | **0.00** |
|  | >44 | -2.80 | -7.98 | -2.04 | 0.13 |
| *Bare soil* | **12** | **6.28** | **1.15** | **11.30** | **0.99** |
|  | **28** | **5.32** | **0.40** | **10.15** | **0.98** |
|  | >44 | 0.89 | -4.01 | 5.66 | 0.63 |

**Fig. 2:**
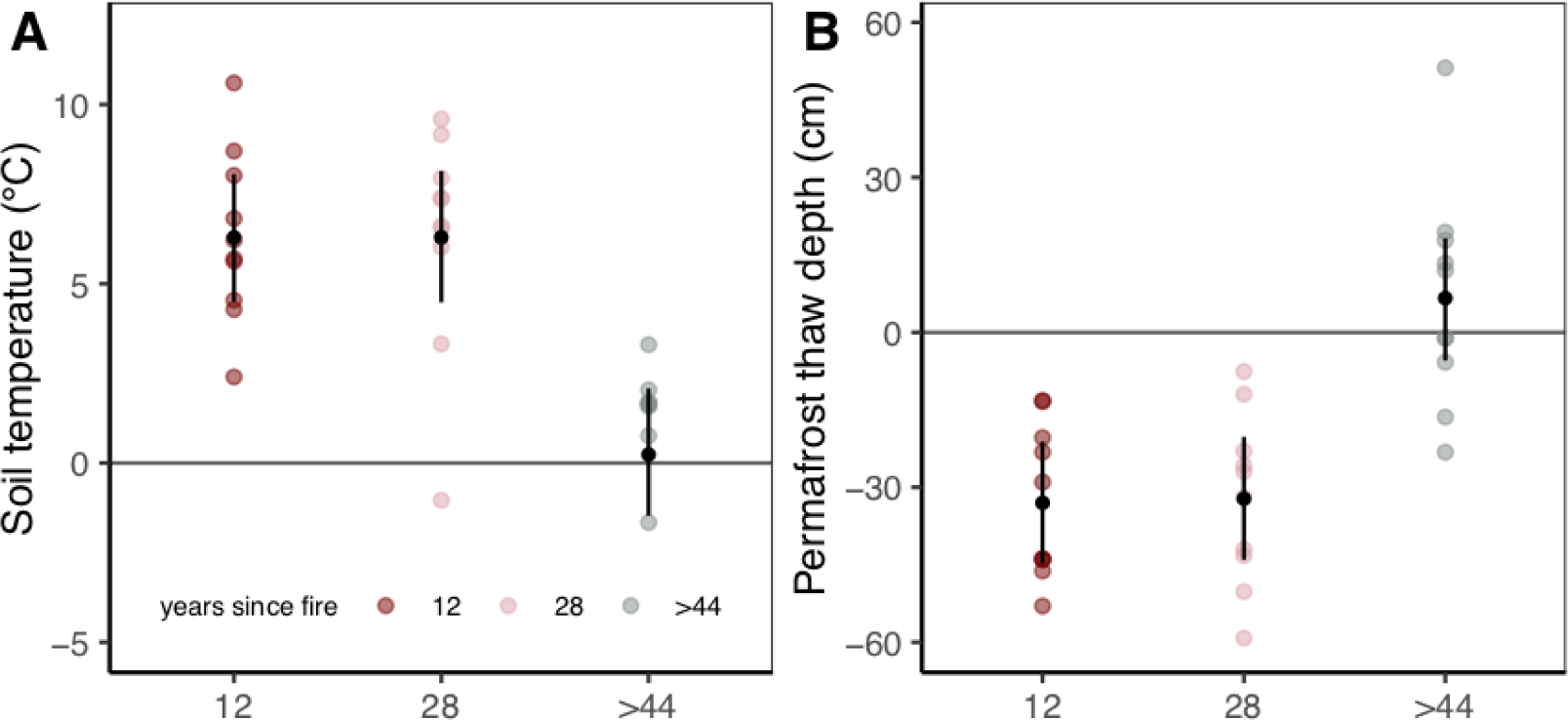
Difference in (A) soil temperature (12 cm depth) and (B) permafrost thaw depth 12, 28 and >44 years after fire. Difference = value on burnt plot minus value on control plot. Coloured dots are differences calculated from the raw data, black dots are predicted mean values and lines 95% credible intervals (CrIs). CrIs not overlapping zero imply a consistent difference between burnt and control plots.

### 3.2. Influence of fire on vegetation cover and height

Bryophyte and lichen cover differed between burnt and unburnt plots and the differences persisted for at least 44 years. On burnt plots, bryophyte cover was 483% higher and lichen cover was 55% lower than on the control plots, more than 44 years after fire (Fig. 3A, Table 2, Suppl. A4).

**Table 2:** Mean values of relative cover and height change after fire, estimated by a linear mixed-effects model, including years since fire and functional groups. Means, 2.5%, 97.5% quantiles of the posterior distributions are given. Effects of change are shown in bold, if there is a high probability of the estimate to be different from 0 (P(*estimate*>0)>0.95 or P(*estimate*>0)<0.05). For the linear mixed-effects model mariginal (marg) and conditional (cond) R^2^ are stated.

| Group | Years since fire | Cover<br>$R^2 = 0.80$ | | | | Height<br>$R^2_{\text{marg}} = 0.19, R^2_{\text{cond}} = 0.23$ | | | |
| --- | --- | --- | --- | --- | --- | --- | --- | --- | --- |
|  |  | Mean | 2.50% | 97.50% | P | Mean | 2.50% | 97.50% | P |
| lichens | 12 | **-42.21** | **-49.85** | **-34.76** | **0.00** | **-7.85** | **-14.44** | **-0.91** | **0.01** |
|  | 28 | **-74.35** | **-82.43** | **-66.66** | **0.00** | **-8.00** | **-14.69** | **-1.47** | **0.01** |
|  | >44 | **-42.00** | **-49.76** | **-34.34** | **0.00** | -3.45 | -10.06 | 3.53 | 0.17 |
| bryophytes | 12 | **8.30** | **0.31** | **16.10** | **0.98** | -0.40 | -7.50 | 6.47 | 0.45 |
|  | 28 | **10.87** | **3.03** | **18.76** | **1.00** | -0.65 | -7.95 | 6.74 | 0.45 |
|  | >44 | **11.93** | **3.75** | **19.35** | **1.00** | 0.70 | -6.05 | 7.44 | 0.58 |
| graminoids | 12 | **6.15** | **-1.86** | **14.08** | **0.95** | **-6.50** | **-12.94** | **0.38** | **0.03** |
|  | 28 | 2.66 | -5.45 | 10.48 | 0.73 | -5.50 | -12.33 | 1.56 | 0.06 |
|  | >44 | 0.27 | -7.11 | 8.10 | 0.51 | -4.00 | -10.27 | 3.00 | 0.13 |
| herbs | 12 | 3.83 | -3.89 | 12.01 | 0.82 | 0.10 | -6.66 | 6.78 | 0.51 |
|  | 28 | 2.57 | -5.37 | 10.28 | 0.75 | 2.17 | -4.90 | 9.04 | 0.74 |
|  | >44 | 0.78 | -6.81 | 8.64 | 0.58 | -5.00 | -11.73 | 1.39 | 0.08 |
| dwarf shrubs | 12 | 1.40 | -6.35 | 9.63 | 0.62 | -2.30 | -9.12 | 4.14 | 0.24 |
|  | 28 | **9.70** | **2.28** | **17.90** | **0.99** | -3.40 | -10.29 | 3.33 | 0.15 |
|  | >44 | **9.30** | **1.10** | **16.71** | **0.99** | 0.00 | -7.09 | 7.09 | 0.49 |
| shrubs | 12 | -5.00 | -12.87 | 2.98 | 0.11 | 1.50 | -5.47 | 8.67 | 0.67 |
|  | 28 | **15.30** | **7.67** | **22.97** | **1.00** | **9.60** | **2.72** | **16.11** | **1.00** |
|  | >44 | **21.30** | **13.68** | **28.84** | **1.00** | **11.00** | **4.45** | **17.96** | **1.00** |

**Fig. 3:**
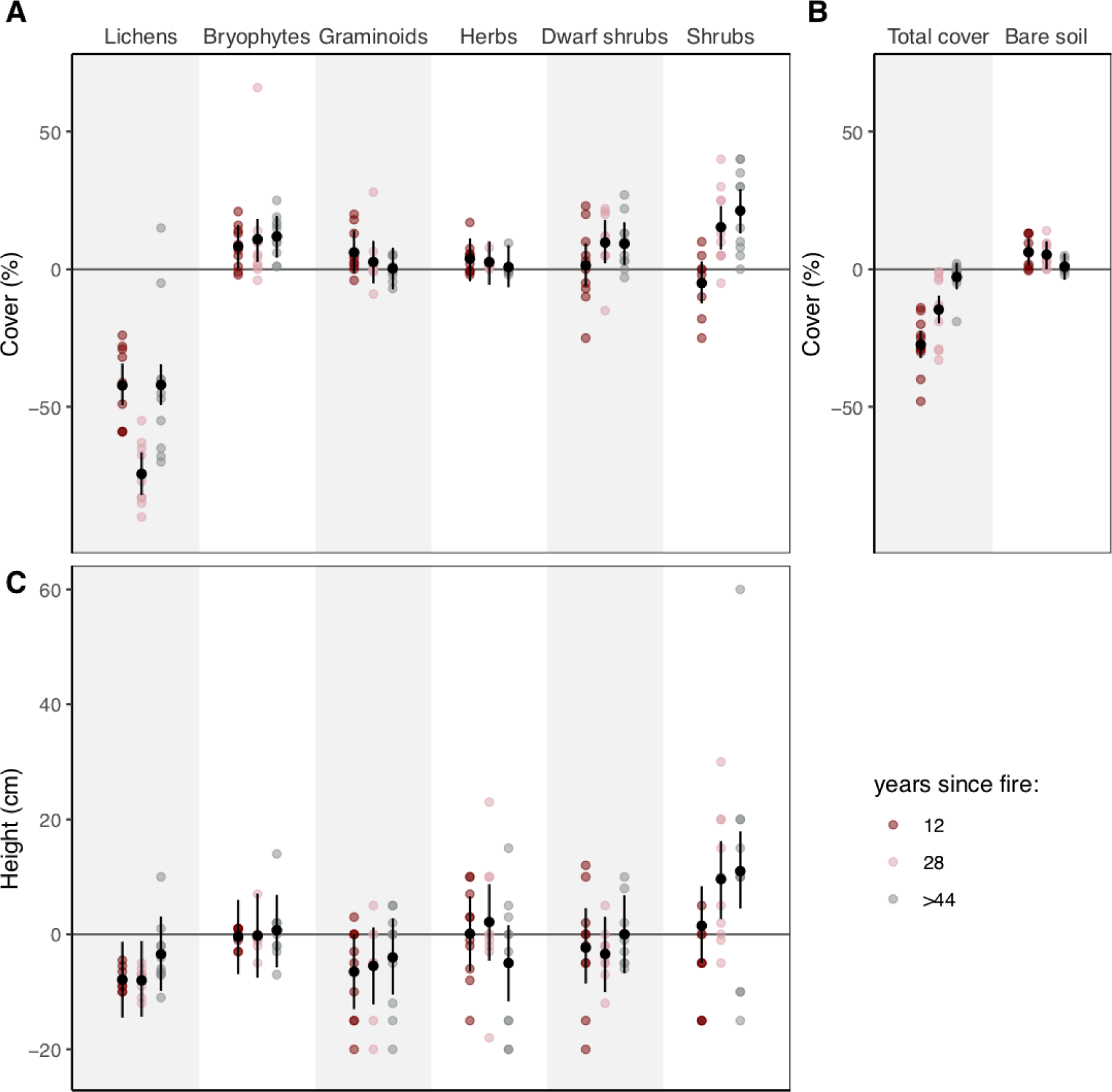
Differences in cover (A) and height (C) for plant functional groups, and differences in total cover and bare soil (B) 12, 28 and >44 years since fire. Difference = value on burnt plot minus value on control plot. Coloured dots are differences calculated from the raw data, black dots are predicted mean values and lines 95% credible intervals CrIs. CrIs not overlapping zero imply a consistent difference between burnt and control plots.

Cover of graminoids was high 12 years after fire but recovered to normal levels within 28 years. Cover of shrubs and dwarf shrubs first did not differ between burnt and unburnt plots, but with increasing time since fire, it increased on the burnt plots. Total cover was lower for 28 years after fire but recovered to control levels after > 44 years. Related to this trajectory decreased bare soil patches after >44 years to normal levels (Fig. 3B, Table 2, Suppl. A4).

Vegetation height was also influenced by fire and the direction of the change was similar as in the vegetation cover (Fig. 3C, Table 2, Suppl. A4): shrubs were taller on sites that burnt 28 and >44 years ago. The height of the lichen carpet was first lower on burnt compared to unburnt plots, but the difference was no longer evident four decades after fire.

### 3.3. Influence of fire on plant traits

Trait values of the dwarf shrub species *Vaccinium uliginosum* did not differ between burnt and unburnt plots. On the other hand, fire had strong and long-lasting impact on traits of the shrub *Betula nana*, in which the mean aboveground cover, height, and SLA were higher on burnt plots (Fig. 4A, B, C and E, respectively, Table 3, Suppl. A5).

**Table 3:**
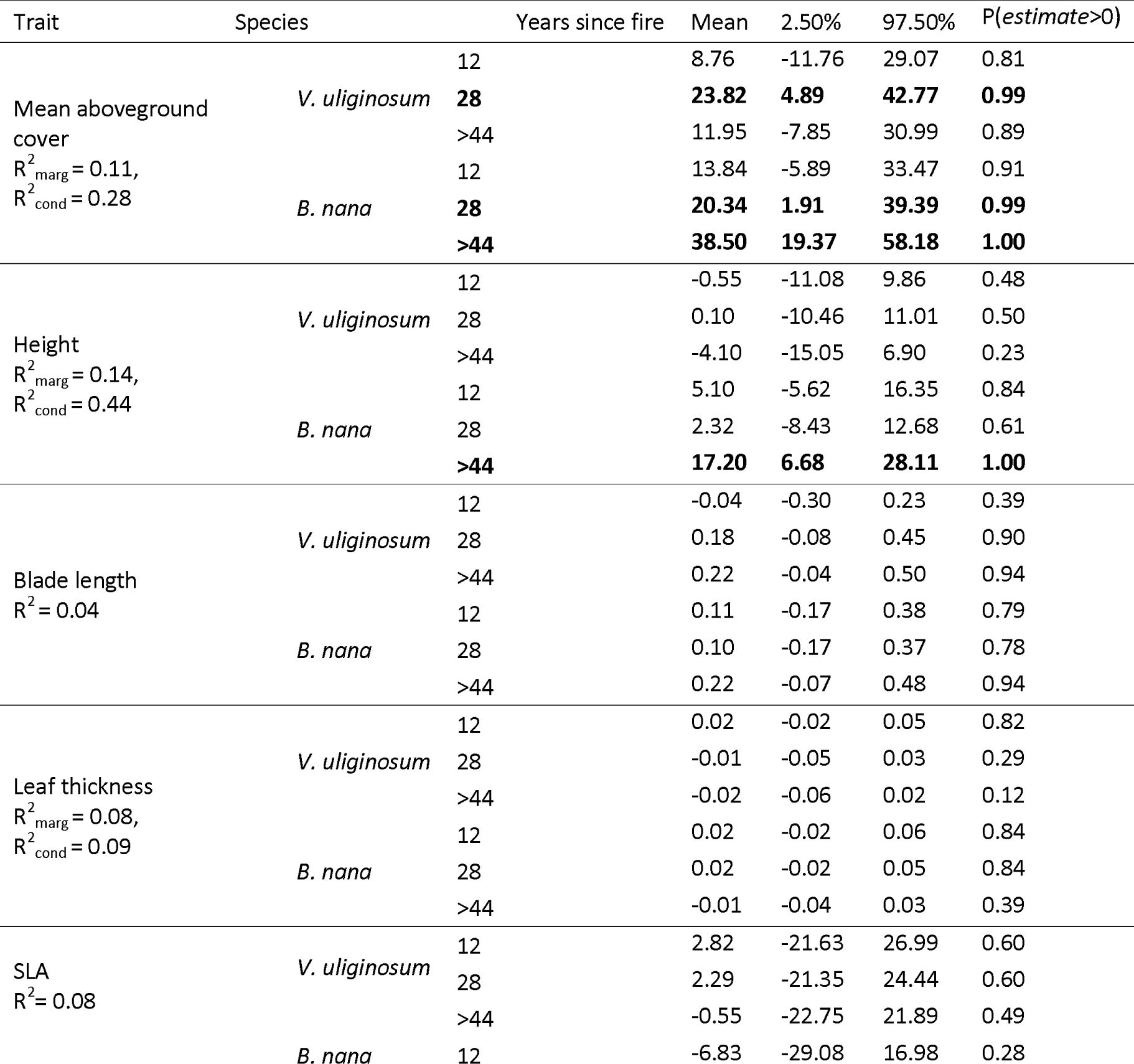
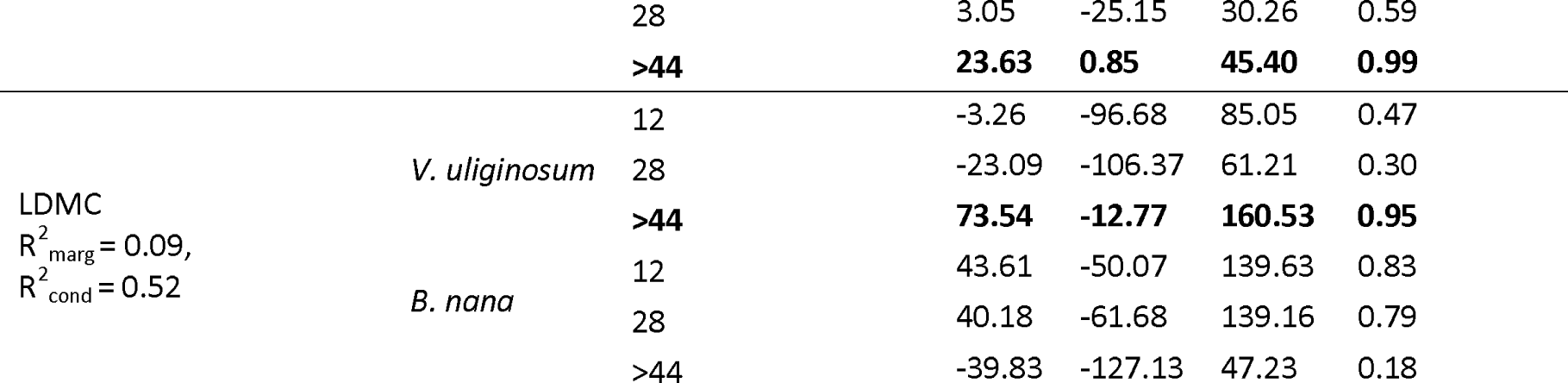
Mean values of change in traits after fire, estimated by linear mixed-effects models, including years since fire and the two different species. Means, 2.5%, 97.5% quantiles of the posterior distributions are given. Effects of change are shown in bold, if there is a high probability of the estimate to be different from 0 (P(*estimate*>0)>0.95). For linear mixed-effects models mariginal (marg) and conditional (cond) R^2^ are stated.

| Trait | Species | Years since fire | Mean | 2.50% | 97.50% | $P(\text{estimate} > 0)$ |
| --- | --- | --- | --- | --- | --- | --- |
| Mean aboveground cover<br>$R^2_{\text{marg}} = 0.11$,<br>$R^2_{\text{cond}} = 0.28$ | *V. uliginosum* | 12 | 8.76 | -11.76 | 29.07 | 0.81 |
|  |  | **28** | **23.82** | **4.89** | **42.77** | **0.99** |
|  |  | >44 | 11.95 | -7.85 | 30.99 | 0.89 |
|  | *B. nana* | 12 | 13.84 | -5.89 | 33.47 | 0.91 |
|  |  | **28** | **20.34** | **1.91** | **39.39** | **0.99** |
|  |  | **>44** | **38.50** | **19.37** | **58.18** | **1.00** |
| Height<br>$R^2_{\text{marg}} = 0.14$,<br>$R^2_{\text{cond}} = 0.44$ | *V. uliginosum* | 12 | -0.55 | -11.08 | 9.86 | 0.48 |
|  |  | 28 | 0.10 | -10.46 | 11.01 | 0.50 |
|  |  | >44 | -4.10 | -15.05 | 6.90 | 0.23 |
|  | *B. nana* | 12 | 5.10 | -5.62 | 16.35 | 0.84 |
|  |  | 28 | 2.32 | -8.43 | 12.68 | 0.61 |
|  |  | **>44** | **17.20** | **6.68** | **28.11** | **1.00** |
| Blade length<br>$R^2 = 0.04$ | *V. uliginosum* | 12 | -0.04 | -0.30 | 0.23 | 0.39 |
|  |  | 28 | 0.18 | -0.08 | 0.45 | 0.90 |
|  |  | >44 | 0.22 | -0.04 | 0.50 | 0.94 |
|  | *B. nana* | 12 | 0.11 | -0.17 | 0.38 | 0.79 |
|  |  | 28 | 0.10 | -0.17 | 0.37 | 0.78 |
|  |  | >44 | 0.22 | -0.07 | 0.48 | 0.94 |
| Leaf thickness<br>$R^2_{\text{marg}} = 0.08$,<br>$R^2_{\text{cond}} = 0.09$ | *V. uliginosum* | 12 | 0.02 | -0.02 | 0.05 | 0.82 |
|  |  | 28 | -0.01 | -0.05 | 0.03 | 0.29 |
|  |  | >44 | -0.02 | -0.06 | 0.02 | 0.12 |
|  | *B. nana* | 12 | 0.02 | -0.02 | 0.06 | 0.84 |
|  |  | 28 | 0.02 | -0.02 | 0.05 | 0.84 |
|  |  | >44 | -0.01 | -0.04 | 0.03 | 0.39 |
| SLA<br>$R^2 = 0.08$ | *V. uliginosum* | 12 | 2.82 | -21.63 | 26.99 | 0.60 |
|  |  | 28 | 2.29 | -21.35 | 24.44 | 0.60 |
|  |  | >44 | -0.55 | -22.75 | 21.89 | 0.49 |
|  | *B. nana* | 12 | -6.83 | -29.08 | 16.98 | 0.28 |
|  |  | 28 | 3.05 | -25.15 | 30.26 | 0.59 |
|  |  | **>44** | **23.63** | **0.85** | **45.40** | **0.99** |
| LDMC<br>$R^2_{\text{marg}} = 0.09,$<br>$R^2_{\text{cond}} = 0.52$ | *V. uliginosum* | 12 | -3.26 | -96.68 | 85.05 | 0.47 |
|  |  | 28 | -23.09 | -106.37 | 61.21 | 0.30 |
|  |  | **>44** | **73.54** | **-12.77** | **160.53** | **0.95** |
|  | *B. nana* | 12 | 43.61 | -50.07 | 139.63 | 0.83 |
|  |  | 28 | 40.18 | -61.68 | 139.16 | 0.79 |
|  |  | **>44** | **-39.83** | **-127.13** | **47.23** | **0.18** |

**Fig. 4:**
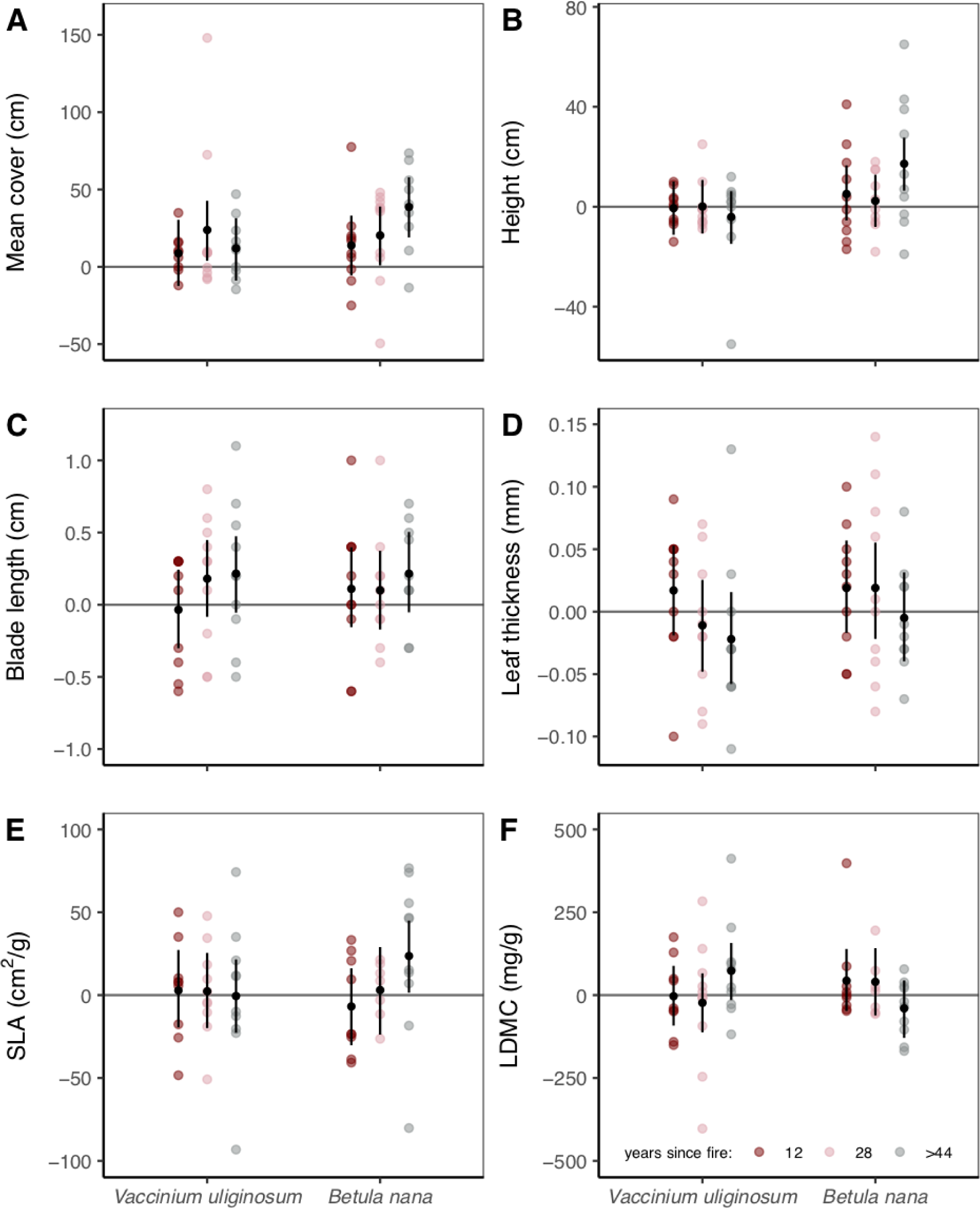
Difference in functional plant trait values of the shrub species *B. nana* and the dwarf shrub *V. uliginosum*, 12, 28 and >44 years after fire. Difference in functional plant trait = plant trait value on burnt plot minus plant trait on control plot. Coloured dots are differences calculated from the raw data, black dots are predicted mean values and lines 95% credible intervals CrIs. CrIs not overlapping zero imply a consistent difference between burnt and control plots.

### 3.4. Vegetation recovery with time since fire

The difference in shrub cover and height between burnt and unburnt plots increased with increasing time since fire. The difference in some functional traits, specifically, aboveground cover and SLA in *Betula nana*, also increased with time. The results were robust regarding uncertainty in the time of the oldest fire, as model results for both scenarios, 44 and 100 years since the last fire, suggested a steep increase of shrub cover and height (Table 4, Table 5).

**Table 4:** Mean slopes of the posterior distribution of relative cover change after fire in regard to time, estimated by linear mixed-effects models for different functional groups. Scenario 1 includes the continuous variable time with raw data related to 12, 28 and 44 years since fire. Scenario 2 includes raw data related to 12, 28 and 100 years since fire. Values are in bold if relative cover change increases strongly with years since fire (P(*slope*>0)>0.95). For linear mixed-effects models mariginal (marg) and conditional (cond) R^2^ are stated.

| Group | Cover |  |  |  | Height |  |  |  |
| --- | --- | --- | --- | --- | --- | --- | --- | --- |
|  | Scenario 1 |  | Scenario 2 |  | Scenario 1 |  | Scenario 2 |  |
| | $R^2_{\text{marg}} = 0.19,$<br>$R^2_{\text{cond}} = 0.22$ | | $R^2_{\text{marg}} = 0.18,$<br>$R^2_{\text{cond}} = 0.22$ | | $R^2_{\text{marg}} = 0.19,$<br>$R^2_{\text{cond}} = 0.22$ | | $R^2_{\text{marg}} = 0.18,$<br>$R^2_{\text{cond}} = 0.22$ | |
|  | Mean | P | Mean | P | Mean | P | Mean | P |
| Lichens | 0.01 | 0.50 | **0.14** | **0.98** | 0.14 | 0.81 | 0.05 | 0.85 |
| Bryophytes | 0.11 | 0.73 | 0.03 | 0.68 | 0.02 | 0.59 | 0.01 | 0.60 |
| Graminoids | -0.18 | 0.16 | -0.06 | 0.19 | 0.07 | 0.68 | 0.02 | 0.69 |
| Herbs | -0.10 | 0.30 | -0.03 | 0.31 | -0.16 | 0.14 | -0.07 | 0.09 |
| Dwarf shrubs | 0.25 | 0.89 | 0.06 | 0.80 | 0.07 | 0.69 | 0.029 | 0.75 |
| Shrubs | **0.82** | **1** | **0.23** | **1** | **0.29** | **0.98** | 0.22 | 0.94 |

**Table 5:** Mean slopes of change within traits of the dwarf shrub species *V. uliginosum* and the shrub species *B. nana* after fire in regard to time, estimated by linear mixed-effects model for different groups. Scenario 1 includes the continuous variable time with raw data related to 12, 28 and 44 years since fire. Scenario 2 includes raw data related to 12, 28 and 100 years since fire. Values are in bold if relative cover change increases strongly (P(*slope*>0)>0.95). For linear mixed-effects models mariginal (marg) and conditional (cond) R^2^ are stated.

| Trait | Species | Scenario 1 |  |  | Scenario 2 |  |  |
| --- | --- | --- | --- | --- | --- | --- | --- |
| | | Mean Slope | $P(\beta>0)$ | $R^2$ | Mean Slope | $P(\beta>0)$ | $R^2$ |
| Mean cover | *V. uliginosum* | 0.09 | 0.58 | marg=0.05 | -0.03 | 0.42 | marg=0.06 |
|  | *B. nana* | **0.80** | **0.97** | cond=0.24 | **0.28** | **0.98** | cond=0.24 |
| Height | *V. uliginosum* | -0.08 | 0.36 | marg=0.12 | -0.04 | 0.31 | marg=0.14 |
|  | *B. nana* | 0.38 | 0.92 | cond=0.39 | **0.16** | **0.97** | cond=0.43 |
| Blade length | *V. uliginosum* | $6.81 \cdot 10^{-3}$ | 0.88 | =0.03 | $1.83 \cdot 10^{-3}$ | 0.80 | =0.03 |
| | *B. nana* | $3.76 \cdot 10^{-3}$ | 0.73 | | $1.84 \cdot 10^{-3}$ | 0.80 | |
| Leaf thickness | *V. uliginosum* | $-1.13 \cdot 10^{-3}$ | 0.09 | =0.07 | $-3.27 \cdot 10^{-4}$ | 0.13 | marg=0.06 |
| | *B. nana* | $-8.13 \cdot 10^{-4}$ | 0.16 | | $-2.49 \cdot 10^{-4}$ | 0.19 | cond=0.08 |
| SLA | *V. uliginosum* | -0.12 | 0.44 | marg=0.06 | -0.04 | 0.42 | =0.07 |
|  | *B. nana* | **0.96** | **0.98** | cond=0.45 | **0.33** | **0.97** |  |
| LDMC | *V. uliginosum* | 2.51 | 0.89 | marg=0.06 | **1.04** | **0.94** | marg=0.08 |
|  | *B. nana* | -2.66 | 0.10 | cond=0.45 | 0.98 | 0.07 | cond=0.51 |

These findings based on the space-for-time substitution approach, were further in agreement with the NDVI trend at the fire scar from 1990. Directly after the fire in August 1990, the NDVI was 40% lower on burnt than on unburnt plots but recovered rather quickly with time (Fig. 5, Suppl. A6). The mean estimate of the posterior distribution was −0.15, slope 5.97*10^−5^ and the quadratic term 3.24*10^−9^ (R^2^ =0.71). After 8 years, NDVI reached the level of unburnt plots and showed a continuous increase thereafter. NDVI values on burnt plots were 26% higher in comparison to unburnt plots, 27 years after fire.

**Fig. 5:**
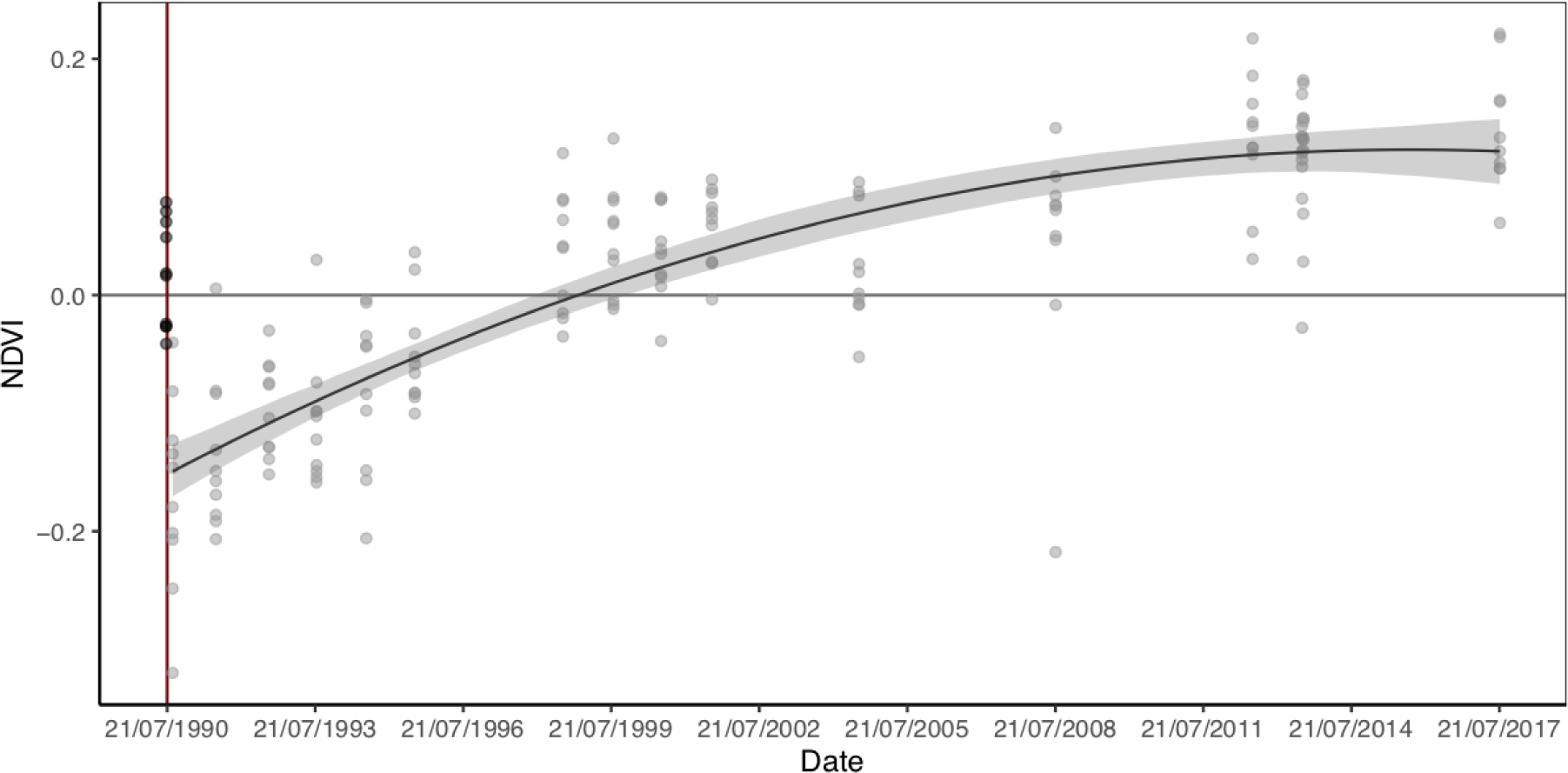
Difference in NDVI (NDVI on burnt plot minus NDVI on control plot) with time. Dots are differences calculated from the raw data of the area which burnt in 1990 and control plots. Black dots indicate values before fire and grey dots after fire. A linear model (LM) (fitted to the data collected after the fire event) is shown as black line and 95% CrIs in grey. The Red line shows the time of the fire event.

## 4. Discussion

In contrast to soil temperature and permafrost thaw depth, vegetation did not recover to the pre-fire state after >44 years. Burnt areas had lower lichen and higher bryophyte and shrub cover, and the dominating shrub species, *Betula nana*, showed higher vitality.

### 4.1. Long-term fire effects on soil temperature and permafrost thaw depth

Fire had strong effects on soil temperature and permafrost thaw depth that persisted for at least three decades. So far, it was rather unclear how rapidly properties of tundra soils return to pre-fire states as previous studies covered relatively short periods of 4-25 years (Mackay 1995, Vavrek et al. 1999, Racine et al. 2004, Rocha and Shaver 2011, Narita et al. 2015). For tussock-shrub tundra, thaw depth was described to reach pre-fire levels after 6 years (Racine et al. 2004). However, the authors of this study described difficulties in detecting changes in the permafrost thaw depth after fire, due to the lack of a control site, and values for the unburnt depths were predicted by a model. Although slight decreases of thaw depth on burnt sites within 3 years after fire were reported (Rocha and Shaver 2011), a literature review showed that permafrost thaw depth was generally deeper in burnt areas in comparison to unburnt areas until 24 years after fire in tundra ecosystems (Rocha et al. 2012). Here we show that the recovery process in dry forest tundra ecosystems can take decades. Even after 28 years, burnt plots had higher soil temperature and soil permafrost thaw depth. The recovery of soil temperature and permafrost thaw depth is probably connected with the recovery of total vegetation cover. Shortly after fire, there is less vegetation and litter that would insulate the soil from heat. The soil surface is furthermore heated by the sun because of the decreased albedo (Rocha et al. 2012, French et al. 2016). Such surface warming leads to permafrost thaw and subsequent increase of permafrost thaw depth and can even cause permafrost degradation (Jones et al. 2015). When vegetation recovers and litter accumulates, soil is again insulated against the atmosphere, which prevents permafrost thawing. As a consequence, soil temperatures as well as permafrost thaw depth can return to pre-fire levels (Michaelides et al. 2018).

### 4.2. Long-term fire influence on vegetation

In contrast to soil temperature, permafrost thaw and total vegetation patterns, plant functional groups did not recover to the pre-fire state. Even four decades after fire, burnt plots had a much lower cover of lichens, but a higher cover of mosses and shrubs.

The long-lasting effects of fire on lichens might be driven by limited re-colonisation abilities. As fire destroys lichens (Jandt et al. 2008), the re-establishment requires dispersal. The most common species in the area, *Cladonia stellaris* and *Cladonia rangiferina*, disperse mainly through thallus fragments (Webb 1998) and, unless transferred by animals, dispersal distances are relatively short (Heinken 1999). While lichens were limited in reaching pre-burn cover in our study, once established they grew generally well in regard to thallus height on the burned sites and after four decades, lichen height on burnt plots reached nearly same mean values as on control plots. This is in line with the findings of Abudulmanova and Ektova (2015) who observed an increasing growth rate of *Cladonia* species with increasing time after fire. Negative fire impacts on lichens are expected to interact with climate change effects. As climate change and fires induce favourable conditions for vascular plants and change fire frequency, total lichen cover in Arctic regions will likely be reduced (Jandt et al. 2008, Joly et al. 2009).

In contrast to lichens, bryophyte cover recovered quickly on burnt plots and even gained higher values than unburnt plots. Bryophytes are often the first plants that colonize burnt areas (Ryömä and Laaka-Lindberg 2005), possibly because they survive in refugia or colonize the area via aerial dispersal (Ryömä and Laaka-Lindberg 2005, Hylander and Johnson 2010). In tundra ecosystems, bryophytes take over the bare ground after fire (Ryömä and Laaka-Lindberg 2005), but they may later compete with herbs and graminoids and decline (Racine et al., 1987). In our study, the cover of bryophytes did not decrease with time since fire, possibly because graminoids were not dominant in the rather dry type of tundra we studied. We did not find any long-term post-fire effect on the cover and height of graminoids and herbs. This contrasts with a number of studies from Alaska that report an increased cover of these functional types after a tundra fire event (e.g. Jones et al. 2013, Narita et al. 2015). One possible reason for this discrepancy might be the time scale. An increased cover of graminoids in the first years was reported by Narita et al. (2015) and Racine et al. (1987), but the studies included only 5-10 years after fire. Even after >100 years, graminoid cover was found to be greater on burnt than on unburnt plots, although graminoid cover was reduced with time since fire (Jones et al. 2013). We did find a slight increase in graminoid height twelve years after the fire, but the differences between burnt and control plots later disappeared, a pattern, that is probably linked to enhanced nutrient availability direct after fire (Jiang et al. 2015b). Another reason for this discrepancy may be vegetation type. While in Alaska, post-fire studies come from moist, tussock tundra types dominated by sedges (*Carex*) and Cottongrass (*Eriophorum*) (Racine et al. 1987, Narita et al. 2015), we studied dry, subarctic tundra with continental climate dominated by lichens.

Our results support findings from other subarctic regions on enhanced shrub growth after fire (e.g. Higuera et al. 2008), which can be evident even >100 years after fire (Jones et al. 2013). Shrubs benefited from fire in our study, which became evident with a considerable delay. Twelve years after fire, shrub cover and height did not differ between burnt and control plots. In the older fire scar (28 years) shrub cover and height increased, and this increase was even more pronounced in the oldest fire scar (>44 years).

In our study, the multitemporal analysis of NDVI in the 1990 fire scar corroborated this trajectory. NDVI analysis showed, that after eight years since fire, greenness approached the level of the unburnt reference and after 22 years a greenness level that is ca. 25 % higher than the reference. The observed increase in NDVI is clearly attributed to the spread of shrubs as the only significantly increasing vegetation component. Our findings are in line with other studies showing that arctic shrubs benefit from fire (Racine et al. 2004, Jandt et al. 2008), reinforcing shrub encroachment (besides other global change related processes) and thereby significantly changing ecosystem functioning (Myers-Smith et al. 2011). This is in contrast to other ecosystems such as tropical savannahs or temperate grasslands where fire is usually counteracting shrub expansion (Naito and Cairns 2011).

Fire can enhance shrub encroachment by two mechanisms. First, fire clears vegetation and thus, shrubs can better germinate and establish (Gough 2006). Second, fire promotes re-growth of shrubs from parts that survived the fire, for example rhizomes (de Groot and Wein 2004). While we do not have any information on seedling establishment, we documented a positive fire impact on growth-related traits, which indicates enhanced growth. Specifically, individual plant cover and height as well as SLA in *B. nana*, the dominating vascular plant species of the shrub tundra in our study region, increased. A similar pattern was shown for alder shrubs (*Alnus viridis subsp. fruticosa*) at the border between subarctic and Arctic (Lantz et al. 2010).

The positive effect of fire on shrub growth is likely mediated via increased surface and soil temperature, availability of nutrients and other ecological factors that change after fire (Chapin 1983, Chambers et al. 2005, Myers-Smith et al. 2015). High surface and soil temperatures generally enhance photosynthesis, nutrient absorption and prolong the vegetation period (Chapin 1983, Nielsen et al. 2017). Higher soil temperatures and deeper permafrost thaw additionally enhance growth by better nutrition through increased rooting depth and soil volume and the provision of surplus nutrients through the stimulated mineralization of organic matter (Rustad et al. 2001, Schuur et al. 2009). While these factors affect all plants, *B. nana* poses further mechanisms that allow the species to benefit from post-fire elevated temperatures. This species can modify its physiology and develop bigger vessels that allow more efficient water transport (Nielsen et al. 2017), and, in contrast to other tundra plants, uses mycorrhiza to transfer belowground carbon (Deslippe and Simard 2011). Furthermore, *B. nana* has the ability to change its growth strategy – with enhanced nutrient availability more long shoots are produced (Bret-Harte et al. 2001). As a result, *B. nana* showed increased height and raised SLA, indicating improved thermal growth conditions and nutrient availability (Kummerow et al. 1987, Shaver et al. 2001) on burnt plots. This relationship between *B. nana* and mycorrhizal networks might be the reason, why we found diverging patterns in the subshrub *V. uliginosum* and *B. nana* in response to fires. The enhanced shrub growth and increased nutrient availability (as indicated by SLA) persisted even in the oldest successional stage (>44 years), although the permafrost thaw depth and soil temperatures returned to similar levels as in unburnt plots. This illustrates a strong fire legacy effect in shrub growth, possibly because individual shrub plants on burnt plots gained growth advantage over shrub plants on control plots during the time-limited phase when soil temperatures were higher and permafrost thaw deeper. The growth advantage remains also after the soil thermal regime went back to normal levels.

## 5. Conclusions

While soil thermal properties and total vegetation cover seemed to recover within decades in our study, recovery of plant functional groups, was not complete after 44 years and it is unclear whether it will recover to a pre-fire state at all. A clear winner of tundra fires in Western Siberia is the shrub species *B. nana*, which showed enhanced growth of individual plants after burning. While the permafrost thaw depth and soil temperatures returned to levels comparable with unburnt plots after 44 years, we found shrubs to grow further. This indicates a strong fire legacy effect, with far reaching implications for the whole ecosystem. Through higher shrub cover, an alteration in decomposition patterns is probable (McLaren et al. 2017) and the changing fire regime in Arctic regions might be influenced as well. Using a space-for-time approach we could unravel long-term fire impacts on the soil thermal regime and vegetation and show connections between fire effects and plant traits. We could demonstrate that those findings agreed with the time series analysis of satellite images. Our results suggest, that the recovery after fire in a dry and lichen-dominated subarctic tundra is very slow and that the ecosystem does not reach pre-fire conditions even after almost half a century.

Most studies on post-fire dynamics were conducted in moist tundra in Alaska. We demonstrated that the post-fire dynamic of the dry, lichen-dominated tundra in Russia may follow different recovery patterns but also confirm panarctic findings of enhanced shrub encroachment after fire. A deeper understanding of links between ecosystem processes and fire impacts requires more studies across the entire, and until now largely unstudied, northern Eurasian tundra belt.

## Acknowledgements

We would like to thank Yuliya Aminova, Denis Bazyk, Marvin Diek, Bettina Haas, Wieland Heim, Liv Jessen, Valeriya Kalchugina, Olga Konkova, Christian Lampei, Kseniya Maryasicheva, Mikhail Moskovchenko, Alexandr Pechkin, Dora Schilling and Farid Sulkarnaev for help with data collection.

We are grateful to the Department for Science and Innovation of the Yamalo-Nenets Autonomous Okrug and Alexei Titovsky, the Interregional Expedition Center “Arctic” and Elena Nyukina, as well as to the Arctic Research Center of the Yamalo-Nenets Autonomous Okrug and Andrey Lobanov for the great and fruitful cooperation. For the good collaboration, we thank also Andrei Tolstikov at the University of Tyumen.

Many thanks to Valentin Klaus for useful comments on an earlier version of the manuscript.

## Supporting information

**Table A1:** Cover in % of functional plant groups on burnt and unburnt plots of the three fire scars.

| Years since fire | 12 |  | 28 |  | >44 |  |
| --- | --- | --- | --- | --- | --- | --- |
| Burn status | unburnt | burnt | unburnt | burnt | unburnt | burnt |
| Lichens | 44.05 | 1.84 | 85.50 | 11.15 | 76.20 | 34.20 |
| Bryophytes | 3.10 | 11.40 | 1.73 | 12.60 | 2.47 | 14.40 |
| Graminoids | 3.45 | 9.60 | 2.66 | 5.32 | 3.58 | 3.85 |
| Herbs | 1.60 | 5.43 | 0.28 | 2.85 | 1.22 | 2.00 |
| Dwarf shrubs | 16.20 | 17.60 | 9.30 | 19.00 | 7.10 | 16.40 |
| Shrubs | 29.00 | 24.00 | 19.20 | 34.50 | 22.20 | 43.50 |
| Trees | 4.80 | 1.15 | 0.85 | 1.31 | 4.04 | 4.45 |

**Fig. A2:**
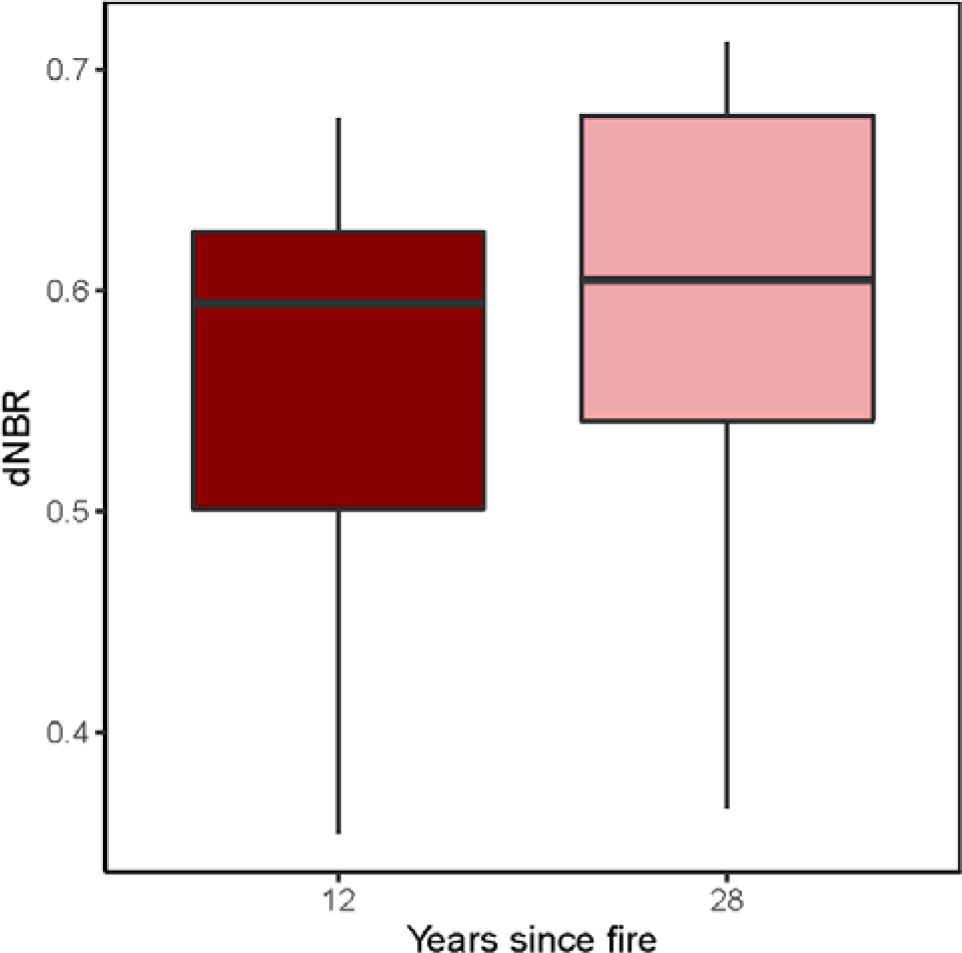
dNBR-values of all burnt plots related to the fire scars burnt in 2005 and 1990.

**Table A3:** Functional groups with names of most abundant species.

| Functional group | Species |
| --- | --- |
| Lichens | *Cladonia stellaris* |
|  | *Cladonia rangiferina* |
| Bryophytes | *Polytrichum juniperinum* |
|  | *Polytrichum commune* |
|  | *Pleurozium schreberi* |
|  | *Aulacomnium turgidum* |
| Graminoids | *Carex globularis* |
|  | *Arctagrostis latifolia* |
|  | *Carex rupestris* |
| Herbs | *Petasites frigidus* |
|  | *Equisetum arvense* |
| Dwarf shrubs | *Vaccinium uliginosum* |
|  | *Vaccinium vitis idaea* |
|  | *Empetrum nigrum* |
| Shrubs | *Betula nana* |
|  | *Ledum palustre* |
|  | *Salix phylicifolia* |
| Trees | *Larix sibirica* |

**Table A4:**
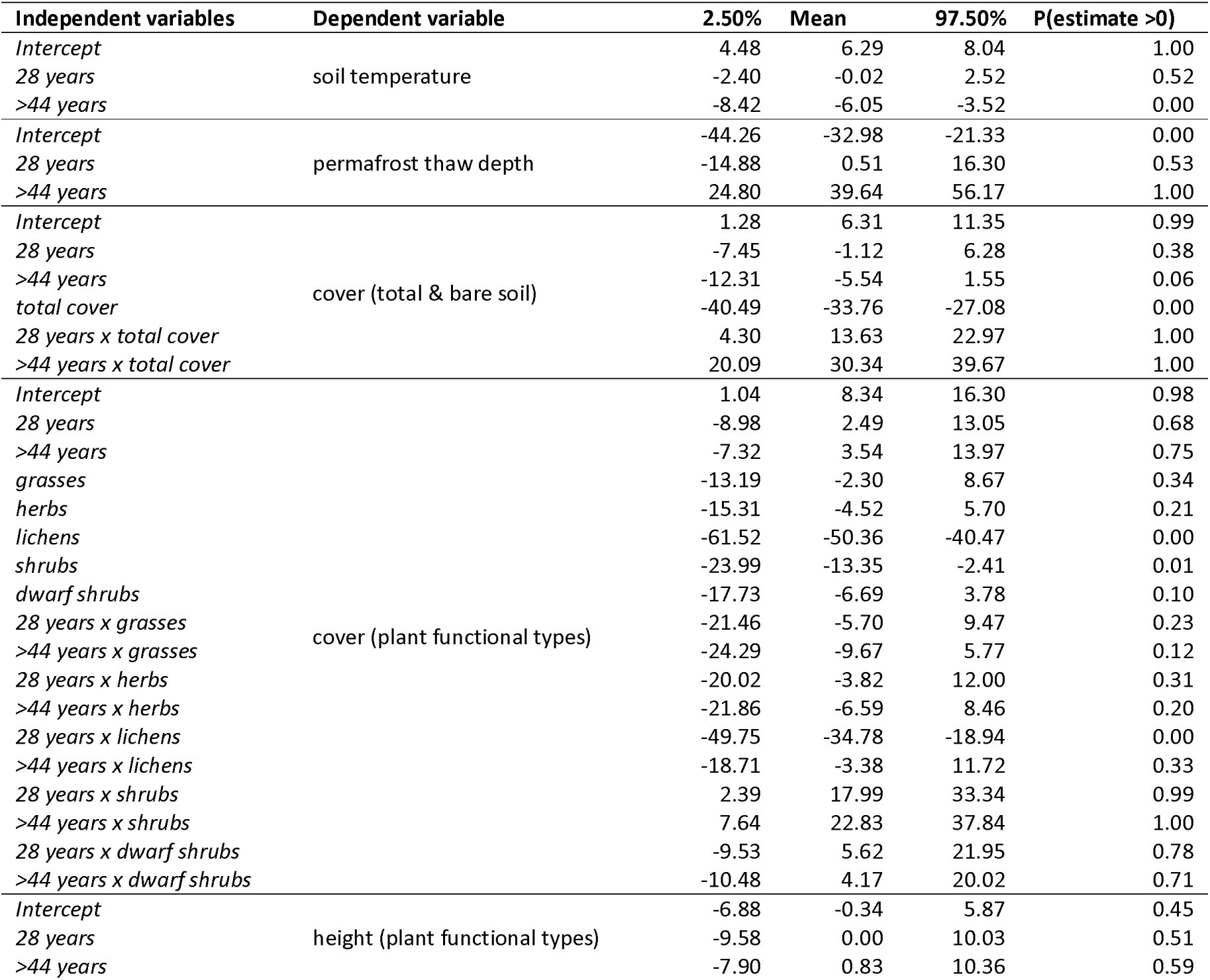
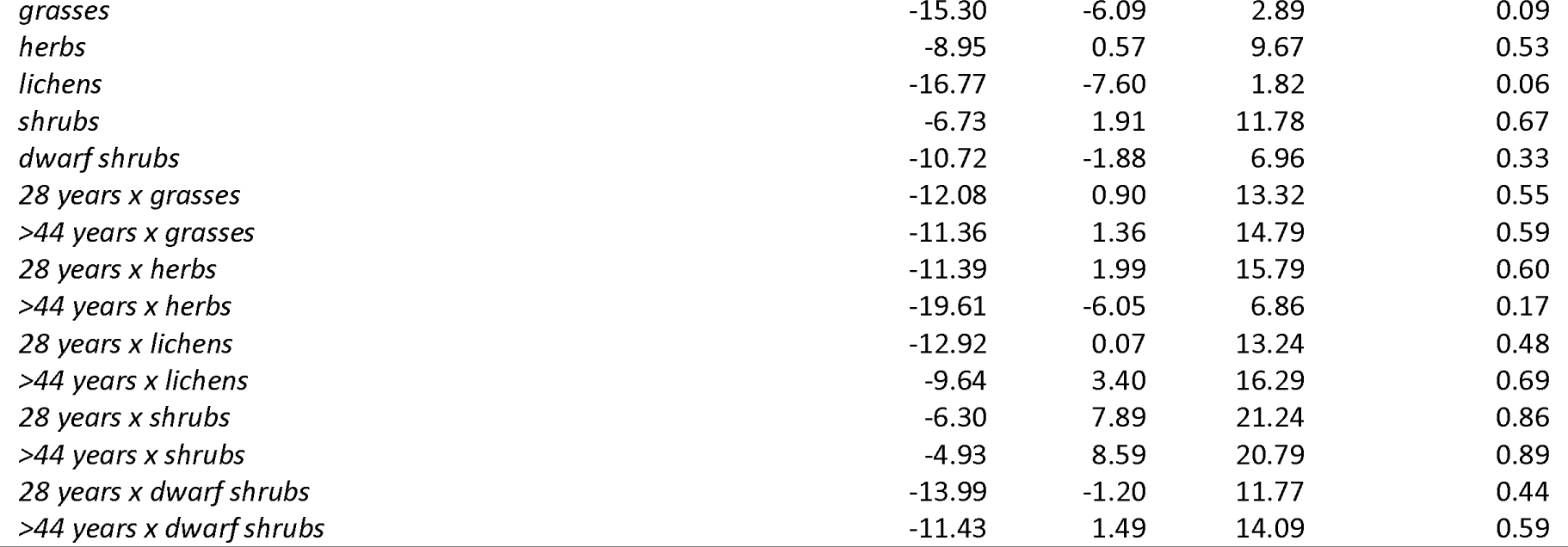
Estimates of linear-mixed effects models of soil temperature, permafrost thaw depth, cover and height in regard to (different groups and) time since fire (12, 28, >44).

| Independent variables | Dependent variable | 2.50% | Mean | 97.50% | P(estimate >0) |
| --- | --- | --- | --- | --- | --- |
| *Intercept* | soil temperature | 4.48 | 6.29 | 8.04 | 1.00 |
| *28 years* |  | -2.40 | -0.02 | 2.52 | 0.52 |
| *>44 years* |  | -8.42 | -6.05 | -3.52 | 0.00 |
| *Intercept* | permafrost thaw depth | -44.26 | -32.98 | -21.33 | 0.00 |
| *28 years* |  | -14.88 | 0.51 | 16.30 | 0.53 |
| *>44 years* |  | 24.80 | 39.64 | 56.17 | 1.00 |
| *Intercept* | cover (total & bare soil) | 1.28 | 6.31 | 11.35 | 0.99 |
| *28 years* |  | -7.45 | -1.12 | 6.28 | 0.38 |
| *>44 years* |  | -12.31 | -5.54 | 1.55 | 0.06 |
| *total cover* |  | -40.49 | -33.76 | -27.08 | 0.00 |
| *28 years x total cover* |  | 4.30 | 13.63 | 22.97 | 1.00 |
| *>44 years x total cover* |  | 20.09 | 30.34 | 39.67 | 1.00 |
| *Intercept* | cover (plant functional types) | 1.04 | 8.34 | 16.30 | 0.98 |
| *28 years* |  | -8.98 | 2.49 | 13.05 | 0.68 |
| *>44 years* |  | -7.32 | 3.54 | 13.97 | 0.75 |
| *grasses* |  | -13.19 | -2.30 | 8.67 | 0.34 |
| *herbs* |  | -15.31 | -4.52 | 5.70 | 0.21 |
| *lichens* |  | -61.52 | -50.36 | -40.47 | 0.00 |
| *shrubs* |  | -23.99 | -13.35 | -2.41 | 0.01 |
| *dwarf shrubs* |  | -17.73 | -6.69 | 3.78 | 0.10 |
| *28 years x grasses* |  | -21.46 | -5.70 | 9.47 | 0.23 |
| *>44 years x grasses* |  | -24.29 | -9.67 | 5.77 | 0.12 |
| *28 years x herbs* |  | -20.02 | -3.82 | 12.00 | 0.31 |
| *>44 years x herbs* |  | -21.86 | -6.59 | 8.46 | 0.20 |
| *28 years x lichens* |  | -49.75 | -34.78 | -18.94 | 0.00 |
| *>44 years x lichens* |  | -18.71 | -3.38 | 11.72 | 0.33 |
| *28 years x shrubs* |  | 2.39 | 17.99 | 33.34 | 0.99 |
| *>44 years x shrubs* |  | 7.64 | 22.83 | 37.84 | 1.00 |
| *28 years x dwarf shrubs* |  | -9.53 | 5.62 | 21.95 | 0.78 |
| *>44 years x dwarf shrubs* |  | -10.48 | 4.17 | 20.02 | 0.71 |
| *Intercept* | height (plant functional types) | -6.88 | -0.34 | 5.87 | 0.45 |
| *28 years* |  | -9.58 | 0.00 | 10.03 | 0.51 |
| *>44 years* |  | -7.90 | 0.83 | 10.36 | 0.59 |

|  |  |  |  |  |
| --- | --- | --- | --- | --- |
| *grasses* | -15.30 | -6.09 | 2.89 | 0.09 |
| *herbs* | -8.95 | 0.57 | 9.67 | 0.53 |
| *lichens* | -16.77 | -7.60 | 1.82 | 0.06 |
| *shrubs* | -6.73 | 1.91 | 11.78 | 0.67 |
| *dwarf shrubs* | -10.72 | -1.88 | 6.96 | 0.33 |
| *28 years x grasses* | -12.08 | 0.90 | 13.32 | 0.55 |
| *>44 years x grasses* | -11.36 | 1.36 | 14.79 | 0.59 |
| *28 years x herbs* | -11.39 | 1.99 | 15.79 | 0.60 |
| *>44 years x herbs* | -19.61 | -6.05 | 6.86 | 0.17 |
| *28 years x lichens* | -12.92 | 0.07 | 13.24 | 0.48 |
| *>44 years x lichens* | -9.64 | 3.40 | 16.29 | 0.69 |
| *28 years x shrubs* | -6.30 | 7.89 | 21.24 | 0.86 |
| *>44 years x shrubs* | -4.93 | 8.59 | 20.79 | 0.89 |
| *28 years x dwarf shrubs* | -13.99 | -1.20 | 11.77 | 0.44 |
| *>44 years x dwarf shrubs* | -11.43 | 1.49 | 14.09 | 0.59 |

**Table A5:**
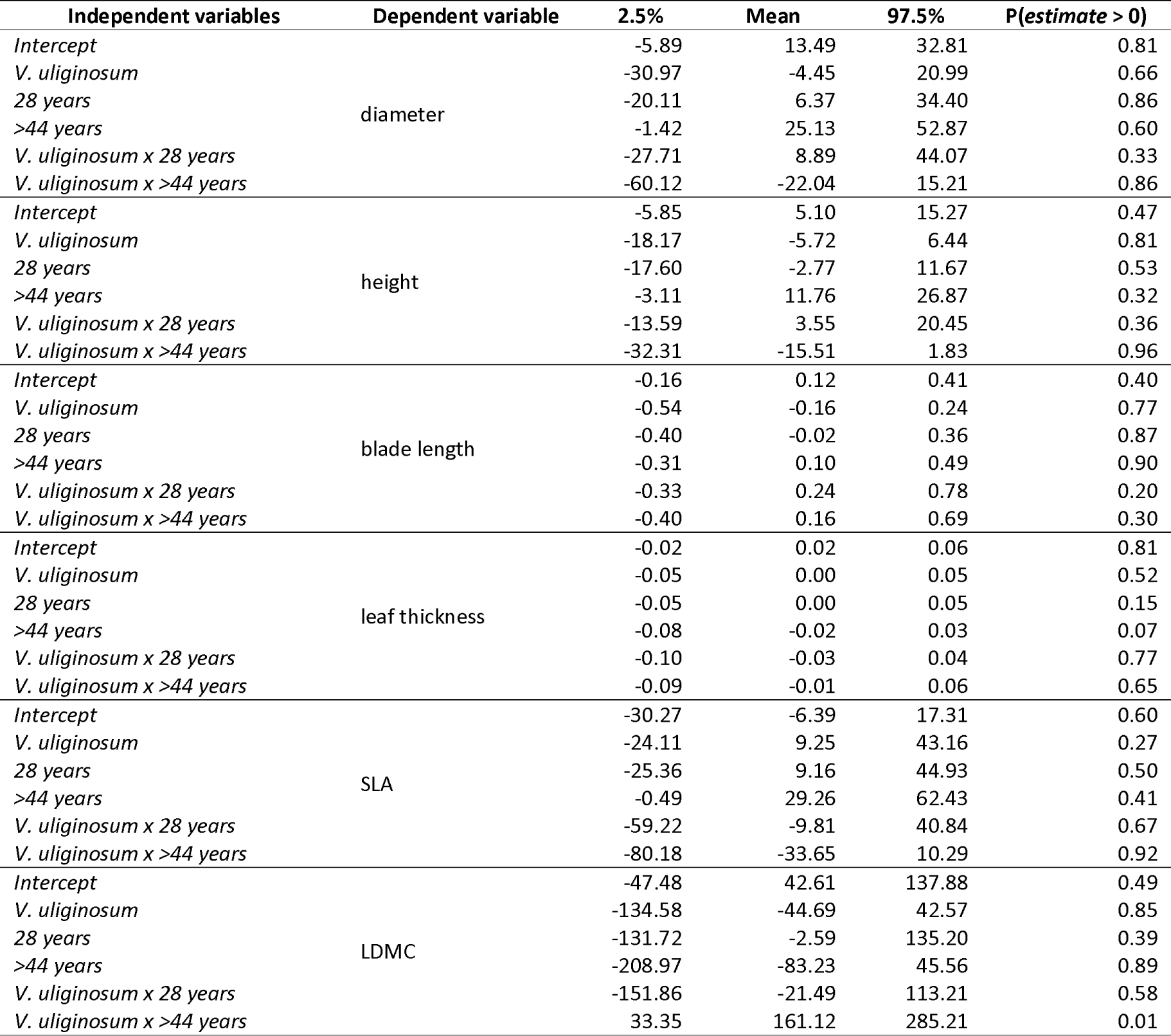
Estimates of linear-mixed effects models of different plant traits in regard to the two species *B. nana* and *V. uliginosum* and time since fire (12, 28, >44).

| Independent variables | Dependent variable | 2.5% | Mean | 97.5% | P(estimate > 0) |
| --- | --- | --- | --- | --- | --- |
| *Intercept* | diameter | -5.89 | 13.49 | 32.81 | 0.81 |
| *V. uliginosum* |  | -30.97 | -4.45 | 20.99 | 0.66 |
| *28 years* |  | -20.11 | 6.37 | 34.40 | 0.86 |
| *>44 years* |  | -1.42 | 25.13 | 52.87 | 0.60 |
| *V. uliginosum x 28 years* |  | -27.71 | 8.89 | 44.07 | 0.33 |
| *V. uliginosum x >44 years* |  | -60.12 | -22.04 | 15.21 | 0.86 |
| *Intercept* | height | -5.85 | 5.10 | 15.27 | 0.47 |
| *V. uliginosum* |  | -18.17 | -5.72 | 6.44 | 0.81 |
| *28 years* |  | -17.60 | -2.77 | 11.67 | 0.53 |
| *>44 years* |  | -3.11 | 11.76 | 26.87 | 0.32 |
| *V. uliginosum x 28 years* |  | -13.59 | 3.55 | 20.45 | 0.36 |
| *V. uliginosum x >44 years* |  | -32.31 | -15.51 | 1.83 | 0.96 |
| *Intercept* | blade length | -0.16 | 0.12 | 0.41 | 0.40 |
| *V. uliginosum* |  | -0.54 | -0.16 | 0.24 | 0.77 |
| *28 years* |  | -0.40 | -0.02 | 0.36 | 0.87 |
| *>44 years* |  | -0.31 | 0.10 | 0.49 | 0.90 |
| *V. uliginosum x 28 years* |  | -0.33 | 0.24 | 0.78 | 0.20 |
| *V. uliginosum x >44 years* |  | -0.40 | 0.16 | 0.69 | 0.30 |
| *Intercept* | leaf thickness | -0.02 | 0.02 | 0.06 | 0.81 |
| *V. uliginosum* |  | -0.05 | 0.00 | 0.05 | 0.52 |
| *28 years* |  | -0.05 | 0.00 | 0.05 | 0.15 |
| *>44 years* |  | -0.08 | -0.02 | 0.03 | 0.07 |
| *V. uliginosum x 28 years* |  | -0.10 | -0.03 | 0.04 | 0.77 |
| *V. uliginosum x >44 years* |  | -0.09 | -0.01 | 0.06 | 0.65 |
| *Intercept* | SLA | -30.27 | -6.39 | 17.31 | 0.60 |
| *V. uliginosum* |  | -24.11 | 9.25 | 43.16 | 0.27 |
| *28 years* |  | -25.36 | 9.16 | 44.93 | 0.50 |
| *>44 years* |  | -0.49 | 29.26 | 62.43 | 0.41 |
| *V. uliginosum x 28 years* |  | -59.22 | -9.81 | 40.84 | 0.67 |
| *V. uliginosum x >44 years* |  | -80.18 | -33.65 | 10.29 | 0.92 |
| *Intercept* | LDMC | -47.48 | 42.61 | 137.88 | 0.49 |
| *V. uliginosum* |  | -134.58 | -44.69 | 42.57 | 0.85 |
| *28 years* |  | -131.72 | -2.59 | 135.20 | 0.39 |
| *>44 years* |  | -208.97 | -83.23 | 45.56 | 0.89 |
| *V. uliginosum x 28 years* |  | -151.86 | -21.49 | 113.21 | 0.58 |
| *V. uliginosum x >44 years* |  | 33.35 | 161.12 | 285.21 | 0.01 |

**Table A6:**
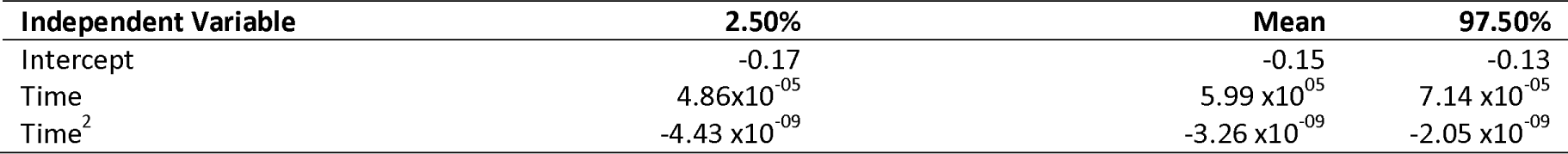
Estimates of linear-mixed effects models for NDVI in regard to time.

## References

Abbott, B. W. et al. 2016. Biomass offsets little or none of permafrost carbon release from soils, streams, and wildfire: An expert assessment. - Environ. Res. Lett. 11: 34014.

Abdulmanova, S. Y. and Ektova, S. N. 2015. Variations in the growth rate of Cladonia lichens during long-term postfire successions in the north of West Siberia. - Contemp. Probl. Ecol. 8: 326–336.

Archibald, S. et al. 2013. Defining pyromes and global syndromes of fire regimes. - Proc. Natl. Acad. Sci. U. S. A. 110: 6445–6447.

Arefiev, S. 2016. The reaction of trees and shrubs in the eastern part of the Tazovsky Peninsula to climate warming. - Ecol. Monit. Biodivers.: 5-9 (in Russian).

Barrett, K. et al. 2012. Vegetation shifts observed in arctic tundra 17 years after fire. - Remote Sens. Lett. 3: 729–736.

Barton, K. 2019. MuMIn: Multi-Model Inference. R package version 1.43.15. https://CRAN.R-project.org/package=MuMIn. in press.

Bates, D. et al. 2015. Fitting Linear Mixed-Effects Models Using lme4. - J. Stat. Softw. 67: 1–48.

Bjorkman, A. D. et al. 2018. Plant functional trait change across a warming tundra biome. - Nature 562: 57.

Bowman, D. M. J. S. et al. 2009. Fire in the earth system. - Science (80-.). 324: 481–484.

Bret-Harte, M. S. et al. 2001. Developmental plasticity allows Betula nana to dominate tundra subjected to an altered environment. - Ecology 82: 18–32.

Bret-Harte, M. S. et al. 2013. The response of Arctic vegetation and soils following an unusually severe tundra fire. - Philos. Trans. R. Soc. Lond. B. Biol. Sci. 368: 20120490.

Chambers, S. D. et al. 2005. Fire effects on net radiation and energy partitioning: Contrasting responses of tundra and boreal forest ecosystems. - J. Geophys. Res. D Atmos. 110: 1–9.

Chapin, F. S. 1983. Direct and indirect effects of temperature on arctic plants. - Polar Biol. 2: 47–52.

Cornelissen, J. H. C. and Makoto, K. 2014. Winter climate change, plant traits and nutrient and carbon cycling in cold biomes. - Ecol. Res. 29: 517–527.

Dahl, E. 1975. Stability of tundra ecosystems in Fennoscandia. - In: Fennoscandian tundra ecosystems. Springer, pp. 231–236.

de Groot, W. J. and Wein, R. W. 2004. Effects of fire severity and season of burn on Betula glandulosa growth dynamics. - Int. J. Wildl. Fire 13: 287–295.

Deslippe, J. R. and Simard, S. W. 2011. Below-ground carbon transfer among Betula nana may increase with warming in Arctic tundra. - New Phytol. 192: 689–698.

Flannigan, M. D. et al. 2009. Implications of changing climate for global wildland fire. - Int. J. Wildl. fire 18: 483–507.

Forbes, B. C. and Kumpula, T. 2009. The ecological role and geography of reindeer (rangifer tarandus) in Northern Eurasia. - Geogr. Compass 3: 1356–1380.

French, N. H. F. et al. 2016. Fire disturbance effects on land surface albedo in Alaskan tundra. - J. Geophys. Res. Biogeosciences 121: 841–854.

Frost, G. V and Epstein, H. E. 2014. Tall shrub and tree expansion in Siberian tundra ecotones since the 1960s. - Glob. Chang. Biol. 20: 1264–1277.

Gelman, A. and Su, Y.-S. 2007. arm: Data Analysis Using Regression and Multilevel/Hierarchical Models. R package version 1.10-1. - URL https://CRAN.R-project.org/package=arm

Gorelick, N. et al. 2017. Google Earth Engine: Planetary-scale geospatial analysis for everyone. - Remote Sens. Environ. 202: 18–27.

Gough, L. 2006. Neighbor effects on germination, survival, and growth in two arctic tundra plant communities. - Ecography (Cop.). 29: 44–56.

Heinken, T. 1999. Dispersal patterns of terricolous lichens by thallus fragments. - Lichenol. 31: 603–612.

Higuera, P. E. et al. 2008. Frequent fires in ancient shrub tundra: Implications of paleorecords for arctic environmental change. - PLoS One 3: 1–7.

Hobbie, S. E. et al. 2002. A synthesis: the role of nutrients as constraints on carbon balances in boreal and arctic regions. - Plant Soil 242: 163–170.

Hu, F. S. et al. 2015. Arctic tundra fires: Natural variability and responses to climate change. - Front. Ecol. Environ. 13: 369–377.

Hudson, J. M. G. et al. 2011. Taller and larger: Shifts in Arctic tundra leaf traits after 16 years of experimental warming. - Glob. Chang. Biol. 17: 1013–1021.

Hylander, K. and Johnson, S. 2010. In situ survival of forest bryophytes in small-scale refugia after an intense forest fire. - J. Veg. Sci. 21: 1099–1109.

IUSS Working Group WRB 2015. World Reference Base for Soil Resources 2014, update 2015 International soil classification system for naming soils and creating legends for soil maps. World Soil Resources Reports No. 106. - FAO.

Jandt, R. et al. 2008. Slow Recovery of Lichen on Burned Caribou Winter Range in Alaska Tundra: Potential Influences of Climate Warming and Other Disturbance Factors. - Arctic, Antarct. Alp. Res. 40: 89–95.

Jiang, Y. et al. 2015a. Contrasting soil thermal responses to fire in Alaskan tundra and boreal forest. - J. Geophys. Res. Earth Surf. 120: 363–378.

Jiang, Y. et al. 2015b. Modeling carbon–nutrient interactions during the early recovery of tundra after fire. - Ecol. Appl. 25: 1640–1652.

Johansson, M. et al. 2013. Rapid responses of permafrost and vegetation to experimentally increased snow cover in sub-arctic Sweden. - Environ. Res. Lett. 8: 35025.

Joly, K. et al. 2009. Decrease of lichens in Arctic ecosystems: The role of wildfire, caribou, reindeer, competition and climate in north-western Alaska. - Polar Res. 28: 433–442.

Jones, B. M. et al. 2013. Identification of unrecognized tundra fire events on the north slope of Alaska. - J. Geophys. Res. Biogeosciences 118: 1334–1344.

Jones, B. M. et al. 2015. Recent Arctic tundra fire initiates widespread thermokarst development. - Sci. Rep. 5: 1–13.

Kazakov, K. 2019. Spravochno-informatsionnyy portal “Pogoda i klimat” [Reference information portal “Weather and Climate”]. -[WWW document] URL http://www.pogodaiklimat.ru/msummary.php?m=all&y=all&id=23256. [accessed 2 May 2019].

Keeley, J. E. et al. 2011. Fire as an evolutionary pressure shaping plant traits. - Trends Plant Sci. 16: 406–411.

Keuper, F. et al. 2012. A frozen feast: Thawing permafrost increases plant-available nitrogen in subarctic peatlands. - Glob. Chang. Biol. 18: 1998–2007.

Key, C. H. and Benson, N. C. 2006. Landscape assessment (LA) - Sampling and Analysis Methods. - USDA For. Serv. Gen. Tech. Rep. RMRS-GTR-164-CD

Korner-Nievergelt, F. et al. 2015. Bayesian Data Analysis in Ecology using Linear Models with R, BUGS and Stan. - Elsevier.

Kornienko, S. G. 2018. Cartography of pyrogenic violations of the vegetation cover on the Tazovsky Peninsula with satellite data. - Actual Probl. oil gas 1: 1–11.

Kummerow, J. et al. 1987. Downslope fertilizer movement in arctic tussock tundra. - Ecography (Cop.). 10: 312–319.

Lantz, T. C. et al. 2010. Response of green alder (Alnus viridis subsp. fruticosa) patch dynamics and plant community composition to fire and regional temperature in north-western Canada. - J. Biogeogr. 37: 1597–1610.

Loranty, M. M. et al. 2014. Siberian tundra ecosystem vegetation and carbon stocks four decades after wildfire. - J. Geophys. Res. Biogeosciences 119: 2144–2154.

Loranty, M. M. et al. 2018. Reviews and syntheses: Changing ecosystem influences on soil thermal regimes in northern high-latitude permafrost regions. - Biogeosciences 15: 5287–5313.

Mack, M. C. et al. 2011. Carbon loss from an unprecedented Arctic tundra wildfire. - Nature 475: 489–492.

Mackay, J. R. 1995. Active layer changes (1968 to 1993) following the forest-tundra fire near Inuvik, NWT, Canada. - Arct. Alp. Res. 27: 323–336.

Masrur, A. et al. 2018. Circumpolar spatio-temporal patterns and contributing climatic factors of wildfire activity in the Arctic tundra from 2001–2015. - Environ. Res. Lett. 13: 14019.

McLaren, J. R. et al. 2017. Shrub encroachment in Arctic tundra: Betula nana effects on above-and belowground litter decomposition. - Ecology 98: 1361–1376.

Mekonnen, Z. A. et al. 2019. Expansion of high-latitude deciduous forests driven by interactions between climate warming and fire. - Nat. plants 5: 952–958.

Michaelides, R. et al. 2018. Inference of the impact of wildfire on permafrost and active layer thickness in a discontinuous permafrost region using the remotely sensed active layer thickness (ReSALT) algorithm. - Environ. Res. Lett. in press.

Mollicone, D. et al. 2006. Ecology: Human role in Russian wild fires. - Nature 440:436.

Moritz, M. A. et al. 2012. Climate change and disruptions to global fire activity. – Ecosphere 3:1–22.

Moskovchenko, D. V et al. 2020. Spatiotemporal Analysis of Wildfires in the Forest Tundra of Western Siberia. - Contemp. Probl. Ecol. 13: 193–203.

Myers-Smith, I. H. et al. 2011. Shrub expansion in tundra ecosystems: Dynamics, impacts and research priorities. - Environ. Res. Lett. in press.

Myers-Smith, I. H. et al. 2015. Climate sensitivity of shrub growth across the tundra biome. - Nat. Clim. Chang. 5: 887–891.

Myers-Smith, I. H. and Hik, D. S. 2013. Shrub canopies influence soil temperatures but not nutrient dynamics: an experimental test of tundra snow–shrub interactions. - Ecol. Evol. 3: 3683–3700.

Naito, A. T. and Cairns, D. M. 2011. Patterns and processes of global shrub expansion. - Prog. Phys. Geogr. 35: 423–442.

Narita, K. et al. 2015. Vegetation and Permafrost Thaw Depth 10 Years after a Tundra Fire in 2002, Seward Peninsula, Alaska. - Arctic, Antarct. Alp. Res. 47: 547–559.

Nielsen, S. S. et al. 2017. Xylem anatomical trait variability provides insight on the climate-growth relationship of Betula nana in western Greenland. - Arctic, Antarct. Alp. Res. 49: 359–371.

Perez-Harguindeguy, N. et al. 2016. Corrigendum to: new handbook for standardised measurement of plant functional traits worldwide. - Aust. J. Bot. 64: 715–716.

R Core Team 2019. R: A language and environment for statistical computing. R Foundation for Statistical Computing. - URL https://www.R-project.org/.

Racine, C. H. et al. 1987. Patterns of vegetation recovery after tundra fires in northwestern Alaska, USA. - Arct. Alp. Res. 19: 461–469.

Racine, C. H. et al. 2004. Tundra Fire and Vegetation Change along a Hillslope on the Seward Peninsula, Tundra Fire and Vegetation Change along a Hillslope on the. 36: 1–10.

Rocha, A. V and Shaver, G. R. 2011. Postfire energy exchange in arctic tundra: the importance and climatic implications of burn severity. - Glob. Chang. Biol. 17: 2831–2841.

Rocha, A. V. et al. 2012. The footprint of Alaskan tundra fires during the past half-century: Implications for surface properties and radiative forcing. - Environ. Res. Lett. in press.

Rustad, L. et al. 2001. A meta-analysis of the response of soil respiration, net nitrogen mineralization, and aboveground plant growth to experimental ecosystem warming. - Oecologia 126: 543–562.

Ryömä, R. and Laaka-Lindberg, S. 2005. Bryophyte recolonization on burnt soil and logs. - Scand. J. For. Res. 20: 5–16.

Sandström, P. et al. 2016. On the decline of ground lichen forests in the Swedish boreal landscape: Implications for reindeer husbandry and sustainable forest management. - Ambio 45: 415–429.

Schuur, E. A. G. et al. 2009. The effect of permafrost thaw on old carbon release and net carbon exchange from tundra. - Nature 459: 556–559.

Shaver, G. R. et al. 2001. Species composition interacts with fertilizer to control long-term change in tundra productivity. - Ecology 82: 3163–3181.

Sundqvist, M. K. et al. 2019. Experimental evidence of the long-term effects of reindeer on Arctic vegetation greenness and species richness at a larger landscape scale. - J. Ecol. 107: 2724–2736.

Turetsky, M. R. et al. 2012. The resilience and functional role of moss in boreal and arctic ecosystems. - New Phytol. 196: 49–67.

U.S. Geological Survey 2018. Earth Explorer.

Vavrek, M. C. et al. 1999. Recovery of Productivity and Species Diversity in Tussock Tundra Following Disturbance. - Arctic, Antarct. Alp. Res. 31: 254.

Viereck, L. A. and Schandelmeier, L. A. 1980. Effects of fire in Alaska and adjacent Canada: a literature review. - US Department of the Interior, Bureau of Land Management, Alaska State Office.

Vilchek, G. E. and Bykova, O. Y. 1992. The origin of regional ecological problems within the northern Tyumen Oblast, Russia. - Arct. Alp. Res. 24: 99–107.

Walker, D. A. et al. 2005. The circumpolar Arctic vegetation map. - J. Veg. Sci. 16: 267–282.

Webb, E. T. 1998. Survival, persistence, and regeneration of the reindeer lichens, Cladina stellaris, C. rangiferina, and C. mitis following clearcut logging and forest fire in northwestern Ontario. - Rangifer 18: 41–47.

Wickham, H. 2009. ggplot2: Elegant Graphics for Data Analysis. - Springer.

Wilke, C. O. 2019. cowplot: Streamlined Plot Theme and Plot Annotations for “ggplot2”. R package version 0.9.4. - URL https://CRAN.R-project.org/package=cowplot.

Young, A. M. et al. 2016. Climatic thresholds shape northern high-latitude fire regimes and imply vulnerability to future climate change. - Ecography (Cop.). 39: 1–12.

Yu, Q. et al. 2015. Land cover and land use changes in the oil and gas regions of Northwestern Siberia under changing climatic conditions. - Environ. Res. Lett. 10: 124020.

Yurkovskaya, T. K. 2011. Vegetation map [Karta rastitel’nosti]. - In: Shoba, S. A. et al. (eds), National Atlas of Soils of the Russian Federation [Natsional’nyy atlas pochv Rossiyskoy Federatsii]. Moscow State University, pp. 46–48.

Zhang, Y. et al. 2015. Spatiotemporal impacts of wildfire and climate warming on permafrost across a subarctic region, Canada. - J. Geophys. Res. Earth Surf. 120: 2338–2356.

